# Organisation of the orthobunyavirus tripodal spike and the structural changes induced by low pH and K^+^ during entry

**DOI:** 10.1101/2022.08.11.503604

**Authors:** Samantha Hover, Frank W Charlton, Jan Hellert, John N Barr, Jamel Mankouri, Juan Fontana

**Affiliations:** School of Molecular and Cellular Biology, University of Leeds, LS2 9JT, United Kingdom; Astbury Centre for Structural and Molecular Biology, University of Leeds, LS2 9JT, United Kingdom; Centre for Structural Systems Biology, Leibniz-Institut für Virologie (LIV), Notkestraße 85, 22607 Hamburg, Germany

**Keywords:** Virus, potassium, bunyavirus, endosome, priming, fusion, glycoprotein, cryo-electron tomography, subtomogram averaging

## Abstract

Following internalisation, viruses employ the changing environment of maturing endosomes as cues to promote endosomal escape, a process mediated by viral glycoproteins. Specifically, we previously showed that both high [K^+^] and low pH promote entry of Bunyamwera virus (BUNV), the prototypical bunyavirus. Here, we used sub-tomogram averaging combined with AlphaFold, to generate a pseudo-atomic model of the whole glycoprotein envelope of BUNV. This allowed us to unambiguously locate the Gc fusion domain and its chaperone Gn within the floor domain of the spike. We also confirmed that low pH and high [K^+^] alters the viral glycoproteins, resulting in an activated intermediate state functionally-distinct from the highly ordered ground state, and we localize these changes to the floor domain. Biochemical data suggests that in this intermediate state the viral fusion loops are partially exposed and selectively interact with host cell membranes. Taken together, we reveal new mechanistic understanding of the requirements for virus entry.

## Introduction

The *Bunyavirales* order is the largest collection of enveloped negative-sense RNA viruses with over 500 named isolates. Within this group, members of the *Hantaviridae, Nairoviridae, Phenuiviridae* and *Peribunyaviridae* families possess tri-segmented genomes (1) that minimally encode an RNA-dependent RNA-polymerase, two viral glycoproteins (GPs) Gn and Gc, which form spikes projecting from the viral envelope, and a nucleocapsid (N) protein that encapsidates the genomic segments forming ribonucleoproteins (2). Several members of these families are serious human pathogens, although no vaccines or antiviral therapies to prevent their associated disease have been approved (3,4).

Bunyaviruses enter cells through the endocytic system, releasing their genomes following fusion between their envelope and vesicular membranes, mediated by the Gn/Gc spikes, which together comprise the fusion machinery. The cues that trigger fusion result from endosomal maturation, which might involve a gradual drop in intraluminal pH, changes in concentrations of other ions or alterations to the lipid and protein composition of endosomal membranes (5), dictating when and where fusion occurs. This prevents premature fusion in inappropriate compartments, which would jeopardize virus entry and infection (6).

For bunyaviruses, the events that lead to spike fusogenesis are not completely characterised. Many bunyaviruses require low pH to induce a fusogenic state and, for some, a high H^+^ concentration has been proposed to be sufficient to establish post-fusion conformations *in vitro* and trigger interactions with liposomes (7–9). Working with the model peribunyavirus, Bunyamwera virus (BUNV), and the model nairovirus, Hazara virus (HAZV), we recently showed the current description of bunyavirus entry is over-simplistic. We showed the concentration of K^+^ ions ([K^+^]) in endosomes, which is controlled by cellular K^+^ channels and increases as endosomes mature, peaking in late endosomes (Fig. 1a), is an important cue for BUNV and HAZV entry. By blocking K^+^ channels pharmacologically, we showed that disruption of endosomal K^+^ accumulation impedes bunyavirus infection by preventing endosomal release, and instead viruses were trafficked to lysosomes, where they were degraded (10), a mechanism that was also interlinked with cellular cholesterol levels (10–12). For HAZV, we further showed that treatment with elevated K^+^ alone (pH 7.3) was sufficient to trigger conformational changes in the GP spikes to induce interactions with co-purified vesicles (13). However, the low resolution of our subtomogram average (STA) and the lack of atomic models for HAZV GPs prevented a detailed structural understanding of the effect of K^+^ on the GPs. Of note, it has recently been shown that disruption of the cellular K^+^ gradient using KCl or the K^+^ ionophore valinomycin inhibits infection by the peribunyaviruses La Crosse virus (LACV), keystone virus and Germiston virus; and of the phenuivirus RVFV (14,15), suggesting that K^+^ is broadly required during bunyavirus entry.

**Fig. 1.**
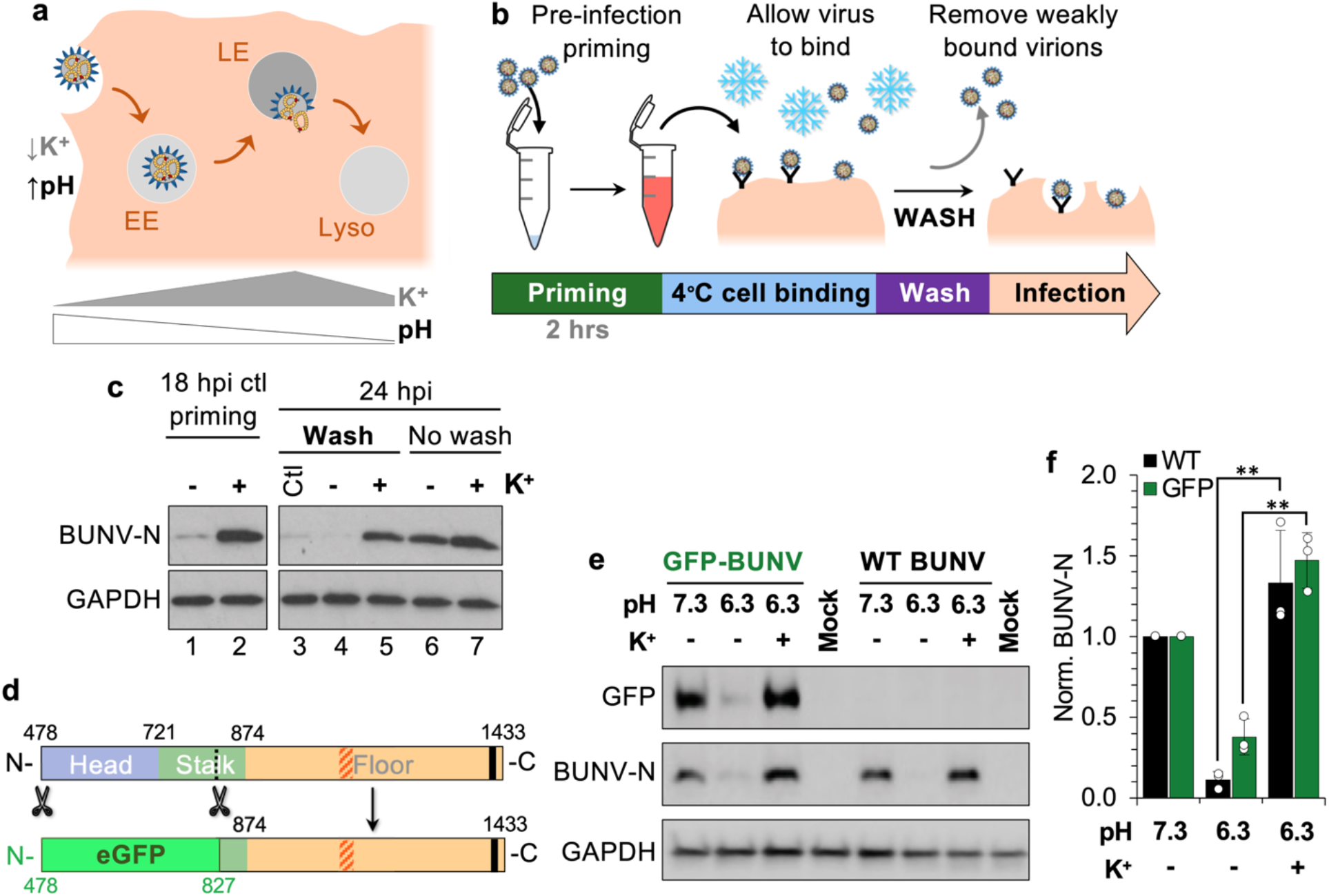
Mimicking endosomal pH and K^+^ in vitro expedites BUNV entry by altering the BUNV GPs. **a** Schematic of the endocytic pathway. The extracellular environment presents a neutral pH (~7.3) and low [K^+^] (~5 mM). Alongside the canonical decrease in pH during endosome maturation, the [K^+^] increases with passage through early endosomes (EE) peaking in late endosome (LE), and is lower within lysosomes (Lyso). The [K^+^] is high in the cytoplasm (~150 mM). (10,54). **b** Workflow of the binding assay, whereby BUNV virions are pre-infection treated for 2 hrs pre-infection with pH 7.3/no K^+^ (Ctl), pH 6.3/no K^+^ or pH 6.3/K^+^ buffers (as previously described (10)). Buffer is diluted out and then virions are bound to cells at 4 °C. Weakly or unbound virions are removed by three PBS washes; or a no wash control. Infection is allowed to proceed for 18 hrs (internal control) or 24 hrs, then cells are lysed and **c** western blot analysed using antibodies against BUNV-N (a known marker of infection) or GAPDH (loading control). **d** Schematic of WT versus eGFP-tagged BUNV Gc protein (GFP-BUNV), with the removed section of the Gc head/stalk domains indicated, which were replaced with eGFP (20). **e** WT and GFP-BUNV were treated with pH 7.3/no K^+^, pH 6.3/no K^+^ or pH 6.3/K^+^, and infection (or non-infected, mock) was assessed by western blot at 18 hpi. **f** Densitometry of n=3 western blots from *d* normalised to each loading control and then to the pH 7.3/no K^+^ control (individual data points shown as white spheres). Error bars indicate mean ± standard error (SE), and significant difference ** = P<0.005 observed between the reduction at pH 6.3/no K^+^ and the recovery at pH 6.3/K^+^.

Here, we used cryo-electron microscopy (cryo-EM) and tomography (cryo-ET) to unravel the structure of the envelope of BUNV and to identify changes in the virion architecture at endosomal pH and [K^+^]. STA was performed to generate an average of the BUNV GPs that allowed us to fit published atomic models of specific regions of Gc (16), and identify the position of the fusion domain of Gc, and of Gn. Modelling both GPs (Gn and Gc) using AlphaFold (17) allowed us to generate a full pseudo-atomic model of the GP envelope of BUNV. Furthermore, we hypothesised that K^+^ triggers BUNV GPs similar to HAZV, causing conformational changes in the distinctive tripodal GP spikes, to elicit a fusion-competent state (13). STA of pH 6.3/K^+^ treated virions revealed drastic changes and an uncoupling of Gn-Gc hetero-hexamers. Consistent with these structural data, biochemical assays confirmed that structural changes elicited by K^+^ are localised to the so-called ‘floor region’ of the GP spikes. pH 6.3/K^+^ treated viruses also interacted more readily with target membranes, suggesting these conditions prompt exposure of the fusion loop as an intermediate step of the GPs prior to fusion triggering.

## Results

### pH 6.3 and K^+^ alter the BUNV GPs to expedite infection

We are interested in the structural and functional changes of bunyavirus GPs in response to biochemical cues during entry. In our previous studies, we showed that the pH and K^+^ dependence of virus entry can be recapitulated *in vitro* (termed ‘priming’), where exposing virions to physiologically relevant endosomal [H^+^] and [K^+^], at 20 mM or higher, led to accelerated onset of infection (10). Here, we aimed to untangle the effects of low pH and K^+^. Of note, when treating viruses at pH 6.3, the addition of K^+^ resulted in a dramatic change in the amount of NP observed at 18 hpi, and suggested a distinct role for K^+^ during entry (10). As the GPs are key mediators of multiple stages of viral entry, we modified the *in vitro* priming assays (10) to assess the effects of K^+^ treatment on GP interactions with host membranes. WT BUNV did not bind strongly to surface receptors at 4 °C (to block endocytosis), with PBS or trypsin washes capable of removing bound virus (Supplementary Fig. 1a). Virions were then treated at pH 6.3/K^+^ (+) or pH 6.3/no K^+^ (-) (or a pH 7.3/no K^+^ treated control (Ctl)), and their binding to host cells assessed. Unbound virions were then removed by washing, and infection was allowed to proceed for 24 h (Fig 1b). A timepoint of 18 hpi was also used to confirm the effects of priming, where K^+^ treated virions would be expected to initiate infection more quickly, showing higher levels of BUNV-N expression (Fig. 1c, lanes 1-2); however, BUNV-N normalises to similar levels as the no-K^+^ treatment by 24 hpi (Fig. 1c, lanes 6-7; as previously described (10)). Importantly, in control and pH 6.3/no K^+^ treated viruses, little-to-no BUNV-N was detectable at 24 hpi when virions were washed after binding at 4 °C, indicating that the washing steps efficiently removed bound virions from the cell surface (Fig. 1c, lanes 3-4). In contrast, BUNV-N was detected at similar levels to the unwashed viruses when virions were pre-treated at pH 6.3/K^+^, indicating that these virions bind more strongly to host cell membranes and/or receptors, and are less readily removed by washing (compare Fig. 1c, lane 5 to lanes 3-4).

We also tested the ability of virions to fuse with the plasma membrane under pH 6.3 conditions using acid-bypass assays (18), using acidic buffers to induce fusion at the plasma membrane and pharmacologically inhibiting endocytic entry (10,12) (Supplementary Fig. 1b). To confirm the effects of pH 6.3/K^+^ during the 2 min of the acid-bypass on viral infection, BUNV-N expression was assessed at 18 hpi and an increase was evident in response to K^+^, albeit at reduced levels, likely owing to the short duration of treatment (2 min versus 2 hrs in ‘priming’ assays above) (Supplementary Fig. 1c). Of note, at 24 hpi, a small BUNV-N band was detected for the bypass at pH 6.3/K^+^ in the presence of drugs that was not present in the no-K^+^ condition, indicating that pH 6.3/K^+^ virions can bypass endocytosis and fuse at the cell membrane. Together, these data indicate that K^+^ at pH 6.3 acts upon the BUNV GPs increasing interactions with target membranes, potentially by exposing the fusion loops.

### K^+^ acts upon the floor region of BUNV GPs

For BUNV, a low resolution structure of the BUNV Gn-Gc (~30 Å) previously revealed an unusual tripod-like structure of the GPs, forming a local lattice-like arrangement of trimers on the virus surface (19). Three regions of the GP tripod were termed the head (apex of the tripod; viral membrane-distal), stalk (connects head-floor) and the floor region (membrane-proximal). More recently, it was identified that the head and stalk regions were formed by the first half of the Gc ectodomain by solving the crystal structures of the head and stalk domains from the peribunyaviruses BUNV (head domain) and Schmallenberg virus (SBV; head-stalk domains) (16). The membrane-proximal floor region is therefore predicted to contain the remaining half of the Gc ectodomain, which includes the fusion domain, and the entire Gn ectodomain.

To identify the region in which K^+^ is acting, we utilised a recombinant BUNV which has the Gc head and ~70% of the stalk replaced with eGFP (GFP-BUNV; Fig. 1d) (20). We hypothesised that if this mutant virus can be triggered by pH 6.3/K^+^ and expedite infection similar to the pH 6.3/K^+^ treated WT BUNV, K^+^ must act on the floor region, as this is the only common component between GPs from WT BUNV and GFP-BUNV. Indeed, this was the case, as evidenced by increased GFP-BUNV-N expression at 18 hpi after pH 6.3/K^+^ treatment (Fig. 1e-f). At this timepoint GFP-BUNV mimicked WT BUNV whereby the presence of K^+^ at pH 6.3 was able to expedite infection significantly compared to the pH 6.3/no K^+^, and to a greater extent than the pH 7.3/no K^+^ control; as previously described for WT (10). This effect on the GFP-BUNV suggests that K^+^ is acting upon the floor region, i.e. Gn and/or the fusion domain of Gc, to increase the efficiency of the fusion machinery. Together with the previous finding that the C-terminal portion of Gc is essential for virus infection (21), this indicates that the floor domain of the envelope is the main determinant for virus entry. However, understanding the architecture and function of the floor region required more structural detail than what was currently available.

### Acidic pH and K^+^ alter the architecture of BUNV virions preventing virion aggregation

To investigate the effects of K^+^ and pH on BUNV GPs by cryo-EM, virus growth and purification conditions were optimised from previously described protocols (19,22). Briefly, BHK-21 cells were infected with BUNV, viral supernatants were collected, concentrated by ultracentrifugation, and titred by plaque assay (Supplementary Fig. 2a-c). SDS-PAGE and negative stain EM of purified BUNV confirmed high viral titres and low background contamination, as required for cryo-EM (Supplementary Fig. 2d,e). To characterise the structural effects of pH and K^+^, purified virions were treated as above at both pH 7.3 and pH 6.3, and with K^+^, prior to vitrification by rapid freezing.

2D cryo-EM micrographs of pH 7.3/no K^+^ treated virions revealed roughly spherical virions in which the lipid bilayer was clearly identifiable surrounding an electron-dense core, likely due to the presence of tightly packed ribonucleoproteins (Fig. 2a; widefield micrographs of cryo-EM images are in Supplementary Fig. 3a-d); as previously described (19). These features were also observed in pH 7.3/K^+^ treated virions (Fig. 2b), though these exhibited a more pleomorphic shape, suggesting structural differences under these conditions or stresses on the virions. At pH 6.3/no K^+^, the morphology of the virions was altered and most virions were forming clusters (Fig. 2c), suggestive of direct interactions between virus particles. In some cases, interacting virions were observed with flattened envelopes adjacent to the interaction site (Fig. 2c, white arrowheads, bottom panels), whereas in other cases, membranes were seen to curve around a neighbouring virion (Fig. 2c, black arrowheads, top panel). We reasoned these interactions may have been driven by exposure of the fusion loops of Gc on one virion contacting neighbouring virions, leading to clustering. Intriguingly, virions at pH 6.3/K^+^ formed no such clusters and appeared as individual virions (Fig. 2d), suggesting that K^+^ prevents the direct interaction of GPs with other viruses thereby preventing clustering at pH 6.3. Taken together, these data suggest that K^+^ has a direct effect on virion morphology and GP architecture and function, beyond that elicited by acidic pH.

**Fig. 2.**
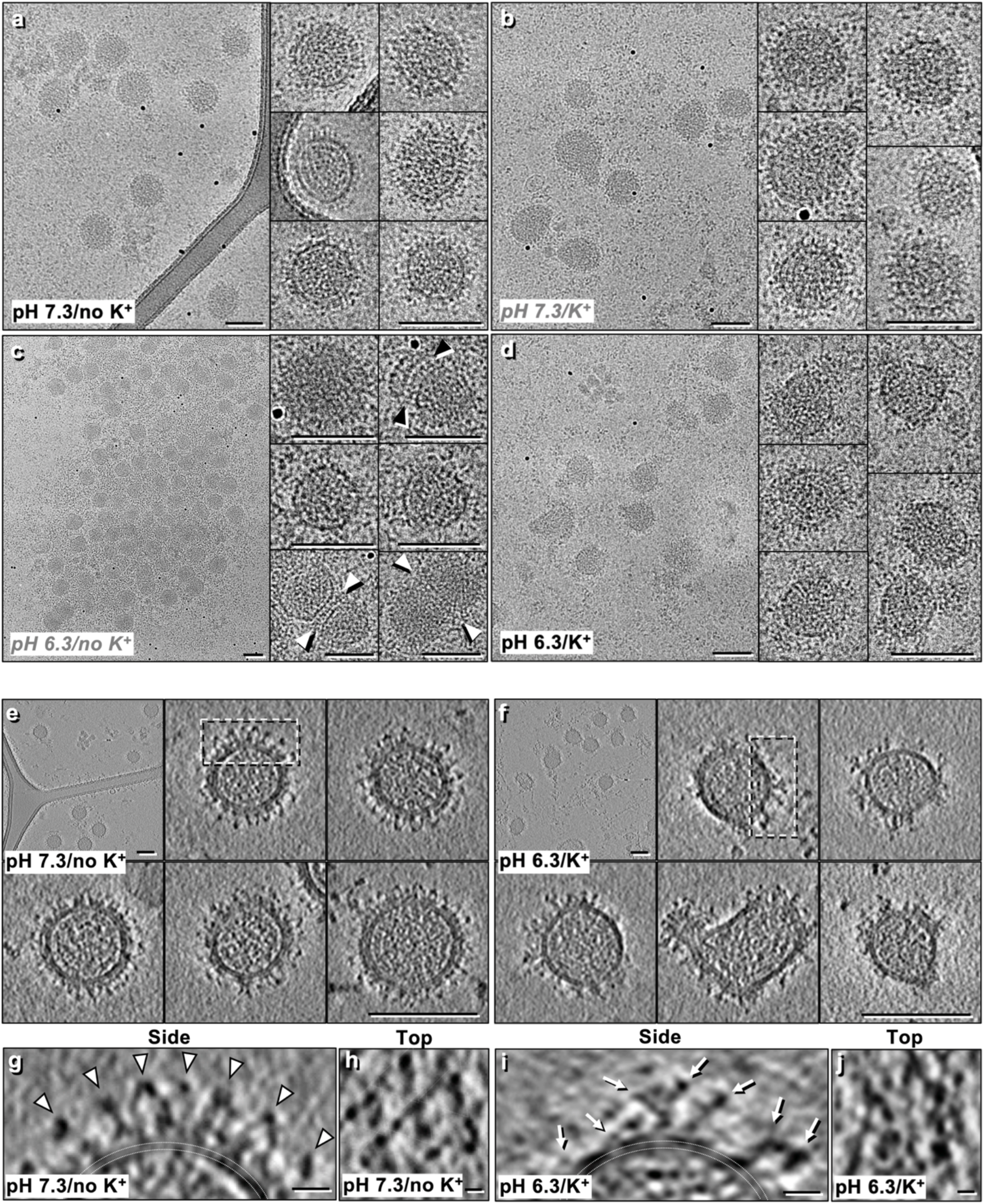
pH 6.3/K^+^ disrupts the BUNV GP spike arrangements on the virion surface. Cryo-EM micrographs of BUNV virions treated under different conditions prior to plunge freezing: **a** pH 7.3/no K^+^, **b** pH 7.3/K^+^, **c** pH 6.3/no K^+^ and **d** pH 6.3/K^+^. White arrowheads in *c* indicate flattened virion contacts with one another and black arrowheads indicate curved membrane contacts. **e-j** Cryo-ET tilt series were collected of pH 7.3/no K^+^ and pH 6.3/K^+^ treated virions, then 3D tomographic reconstructions were calculated. **e** Slices through tomograms of pH 7.3/no K^+^ treated virions (control). **f** As in *e*, with pH 6.3/K^+^ treated virions which show clear disruption to the regular arrangement of GPs. **g,h** Slice through the side (from the white dashed box in *e*) and top of a pH 7.3/no K^+^ treated virion. White arrows indicate tripodal arrangement and triangles in *h* emphasise the trimeric architecture and local lattice. **i,j** As in *g,h*, however from pH 6.3/K^+^ virions where side (from box in *f*) and top views show a disruption to the GP tripods and the local lattice. White arrows indicate spike ‘ends’ that no longer appear connected in tripods. Scale bars: *a-f* = 100 nm; *g-j* = 10 nm.

### Cryo-ET reveals an altered GP arrangement in pH 6.3/K^+^ virions

We next employed cryo-ET, which allows 3D imaging of pleomorphic specimens, to characterise the morphology of virions treated in conditions mimicking the extracellular environment (pH 7.3/no K^+^) and late endosomes (pH 6.3/K^+^) (Fig. 1a) (35). pH 7.3/no K^+^ virions contained an electron-dense core and a continuous arrangement of GPs on the viral surface (Fig. 2e). In contrast, the pH 6.3/K^+^ virions were noticeably more pleomorphic, with clear disruption to the regular GP arrangement (Fig. 2f). Areas of GP-free membrane were also evident on the majority of pH 6.3/K^+^ virions. These data suggested that pH 6.3/K^+^ induced disruption to the local GP arrangements and allows GPs to move more freely and polarise towards one side of the viral membrane and/or that the virus sheds GPs. This polarisation may occur in preparation for interaction with target membranes, leaving large areas of the envelope devoid of GPs.

In our tomograms, both the tripodal structure and local trimeric lattice were easily identifiable on pH 7.3/no K^+^ virions (Fig. 2g,h). However, pH 6.3/K^+^ virions showed a clear disruption of this arrangement and minimal evidence of tripods (Fig. 2i,j), resembling the disruption previously observed in 2D cryo-EM at pH 5.1 (19). Serial sections through individual virions recapitulated these findings; a clear difference was observed in GP architecture and virion morphology between both conditions (Supplementary Fig. 2e,f). Of note, there were no obvious differences in the cores of the virions, suggesting that the effect(s) of pH 6.3/K^+^ were limited to viral GPs. Taken together, these data confirm that treatment at pH 6.3/K^+^ leads to drastic changes in the GP structure of BUNV.

### An improved STA of the BUNV GP permits modelling of the Gc and Gn hetero-hexameric arrangement

To assess the physiological relevance of the changes induced by pH 6.3/K^+^, a higher resolution map of BUNV GP than that available was required. STA was used to align and average pH 7.3/no K^+^ GPs, which confirmed the tripodal arrangement, resolved at ~16 Å (by gold-standard (GS)-FSC Supplementary Fig. 4a). In this average, both leaflets of the viral membrane (‘M’) were clearly defined at the base, with a GP spike projecting ~16 nm from the surface (Fig. 3a,b). This tripod is predicted to constitute a heterohexamer composed of three copies of the smaller Gn (32 kDa, 186 residue ectodomain) and three copies of the larger Gc (110 kDa, 910 residue ectodomain) (23) (Fig. 3c). Our STA showed distinguishable head, stalk and floor regions (Fig. 3d,e), and a previously unidentified density protruding from the floor but shielded by the tripod head (Fig. 3d white arrowheads). Fitting of the previously determined X-ray structures of the head (BUNV) and stalk (SBV) domains into our model (Fig. 3f,g) revealed a tight fit corresponding to cross-correlation values of 0.70 and 0.82 for the head and stalk domains, respectively. The fit of the head trimer allowed us to confirm the handedness of the average (Supplementary Fig. 4b,c). By exclusion, the floor domain must contain the conserved class II fusion domain of Gc and Gn, however specific arrangements could not be obtained from this average (Fig. 3d-f gold region and Supplementary Movie 1).

**Fig. 3.**
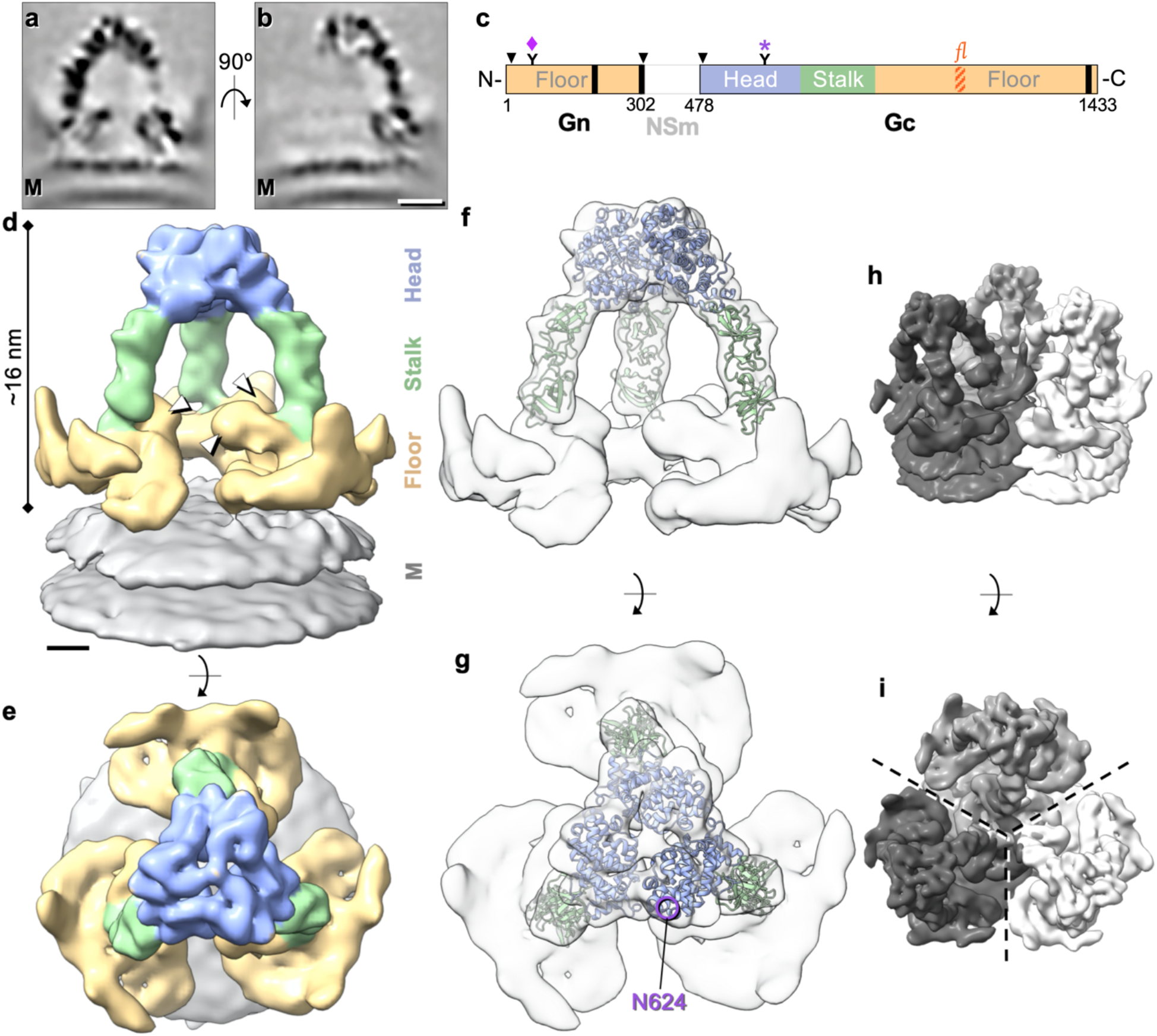
BUNV GPs exhibit a trimeric tripod-like architecture. STA of over 20,000 GP spikes aligned through iterative refinements, generating a ~16 Å average (GS-FSC) (EMD-15557). **a,b** Sections through the electron density averages, showing a trimeric spike on top of the viral lipid bilayer (M; membrane). **c** Schematic of the M segment polyprotein constituted by Gn, NSm and Gc. The Gc protein is comprised of three sections: the head, stalk and floor regions. The fusion loop (*fl;* orange striped) resides in the floor region. Arrowheads indicate cleavage sites, black bars the predicted TMDs, numbers the amino acid residues, and ‘Y’ the glycans N60 (Gn, ♦) and N624 (Gc, *, mAb-742 binding epitope) (16,21,29,30). **d,e** Isosurface rendering of the GP trimer, identifying the three regions; head (light blue), stalk (light green) and floor (gold; where Gn also resides) on top of the viral membrane (grey). White arrowheads indicate a previously unresolved region in the floor. **f,g** The previously solved X-ray structures of the BUNV head (pdb: 6H3V; light blue) and SBV stalk (pdb: 6H3S; light green) domains were fitted into our model (16). The glycosylation site N624 is indicated (purple circle). **h,i** A model of the local lattice arrangement of BUNV GPs, emphasising the two C3 symmetry axes; one forming a tripod, and one in the floor region where three neighbouring tripods connect. Scale bars: *a*, *b* = 5 nm; *d, e* = 2 nm.

To further our understanding of the architecture of this floor domain, STA was performed focused on this region (Fig. 3h,i) and resulted in an average at a slightly improved resolution of ~14 Å, as determined by GS-FSC (Supplementary Fig. 4a). This average is thought to represent a hetero-hexameric assembly of a trimer of Gn and a trimer of the Gc floor region, which includes the fusion domains (Fig. 4a-d). In the floor domain, novel details were identified that were not resolved in the previously published structure (19), highlighting the interconnected organisation of the fusion domains (Fig. 4c,d). Three stalk domains were also resolved, which identify the connections to three independent tripods. A post-fusion structure of the fusion domain for the related peribunyavirus LACV has been recently solved (pdb: 7A57). LACV Gc exhibits a class II fusion domain and shares ~49% amino acid identity and ~69% similarity with the fusion domain of BUNV Gc. We therefore modelled a LACV pre-fusion domain, based on the differences between the pre-fusion and post-fusion conformations observed for other bunyavirus class II fusion domains, which in broad terms involve the rigid-body rearrangement of domain III (Supplementary Fig. 5) (7,24–26). In addition, there was unoccupied density remaining in the floor region that could accommodate a trimer of Gn in the centre, allowing it to contact the Gc fusion domains and the viral membrane (Fig. 4e,f).

**Fig. 4.**
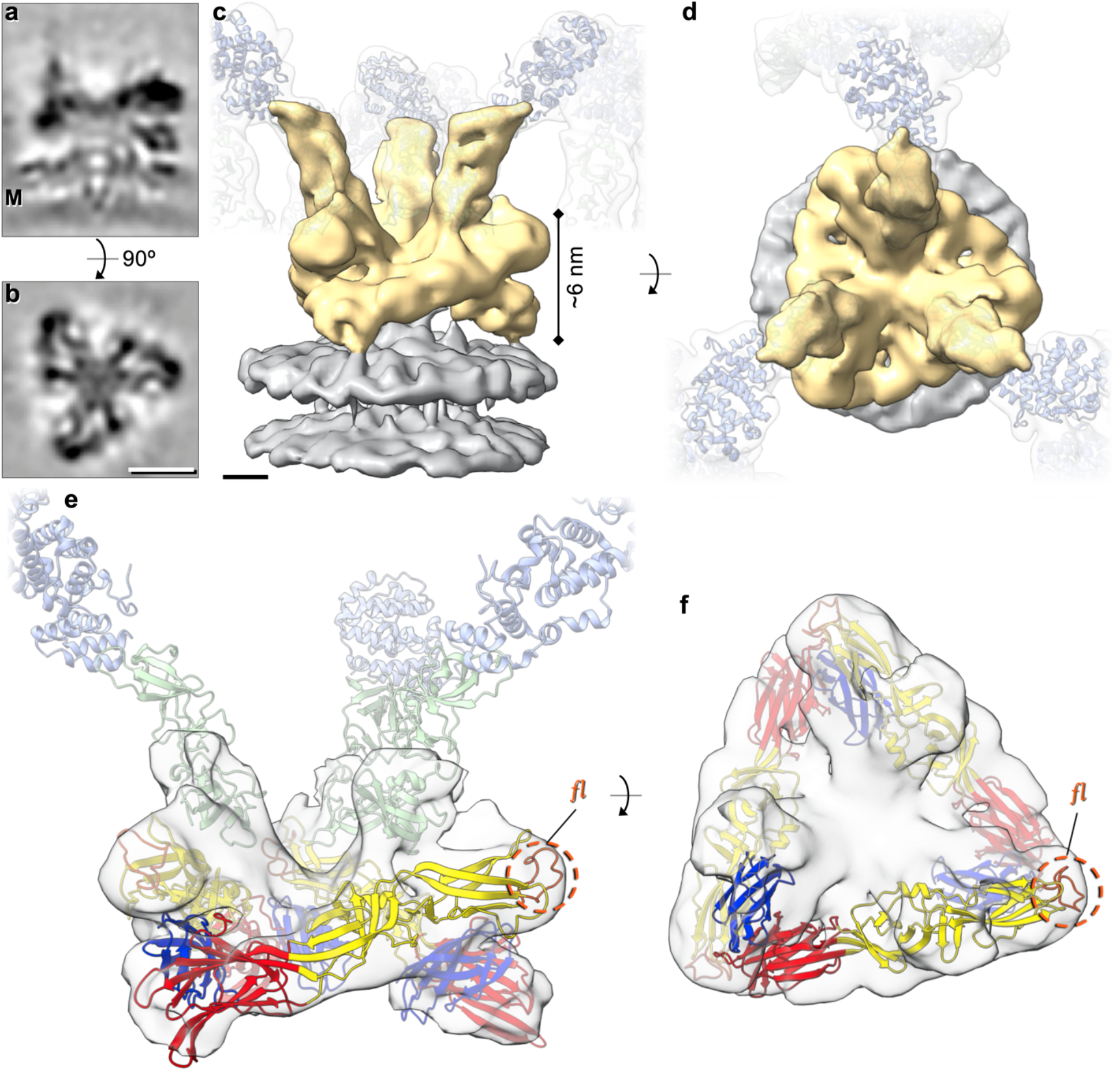
STA of the floor region allows modelling of the BUNV fusion domain. STA was performed as in *Fig. 3*, however aligning the floor region trimer (as opposed to the tripod). Over 16,000 subtomograms were resolved at ~14 Å resolution (GS-FSC). **a,b** Sections through the electron density of the floor region average atop the viral membrane (M). **c,d** Isosurface rendering of the electron density (*c* side view, *d* top view) of the floor region (gold) atop the viral membrane (grey). Translucent tripods of the head-stalk regions have been added for orientation. **e,f** Modelling of the LACV fusion domain (residues 949-1344; domain I red, domain II yellow, domain III blue) in C3 symmetry within the BUNV floor region (EMD-15569), revealing the location of the fusion loops (*fl;* orange) (post-fusion structure pdb: 7A57). The location of the head (light blue) and stalk (light green) domains are indicated in *e* for orientation. Scale bars: *a, b* = 5 nm; c, *d* = 2 nm.

There are currently no Gn structures available for any peribunyavirus and only a head domain structure available for BUNV Gc. In order to interpret the STA map of the BUNV prefusion spike we generated an AlphaFold model of the entire Gn-Gc complex (Fig. 5a,b) (17,27), which reached an overall high confidence score with an average pLDDT of 79 (predicted local distance difference test, which is a per-residue estimate of its confidence; scores of 70-90 are expected to be modelled well (17)) (Supplementary Fig. 6a,b) with all conserved cysteines forming plausible disulphides. Fitting of the ectodomains of Gn and Gc from the AlphaFold model into our STA of the tripod (Fig. 5c,d) required only minor adjustments of the angles between head and stalk, and between the two stalk subdomains (Supplementary Fig. 6c), and it resulted in only minor clashes at protein contact sites within the floor.

**Fig. 5.**
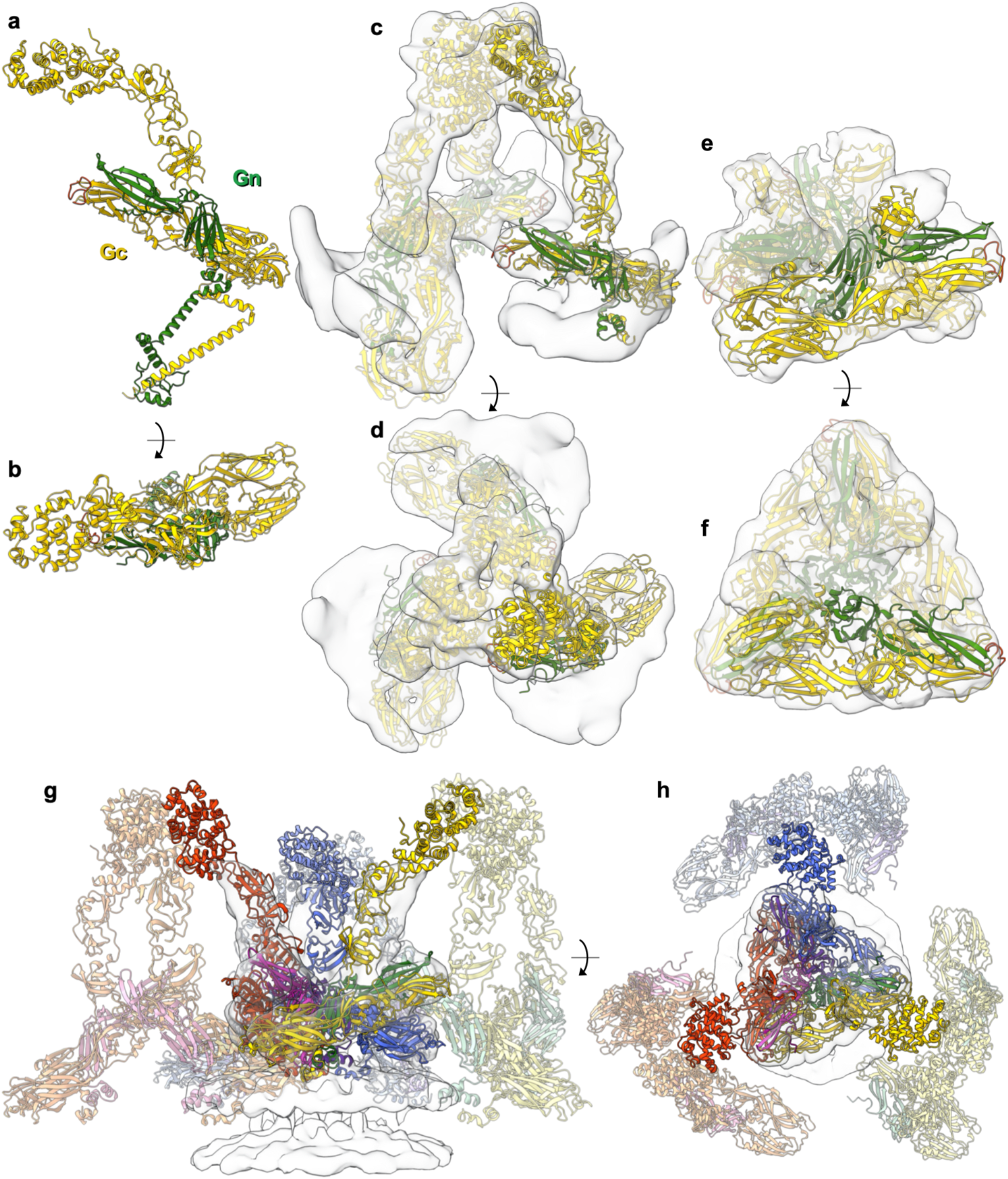
AlphaFold generates a dimeric structure of BUNV Gn and Gc which fits reliably within our GP STA. **a,b** AlphaFold generated an atomic model of the full-length Gn-Gc dimer. Gc in yellow, Gn in green. **c,d** With only minor rotations about the inter-domain regions (*Supplementary Fig. 6c*), this model fitted well within the tripod STA (from *Fig. 3*; AlphaFold pdb found in *Supplementary Material 1*) in C3 symmetry (Gn-Gc ectodomains are shown; residues Gn 1-187, Gc 1-899; no TMD). **e,f** The Gn ectodomain and the Gc fusion domain with the stalk subdomain II (residues 351-899) additionally fitted well within the floor region STA (from *Fig. 4*) in C3 symmetry. **g,h** A model of the BUNV GP envelope can be predicted, here showing three hetero-hexamers fitted within a floor STA. The Gn-Gc pairings are colour coded as follow; hexamer 1 purple-blue, hexamer 2 green-yellow, hexamer 3 pink-red. *Supplementary Movie 2* also demonstrates this envelope model.

**Fig. 6.**
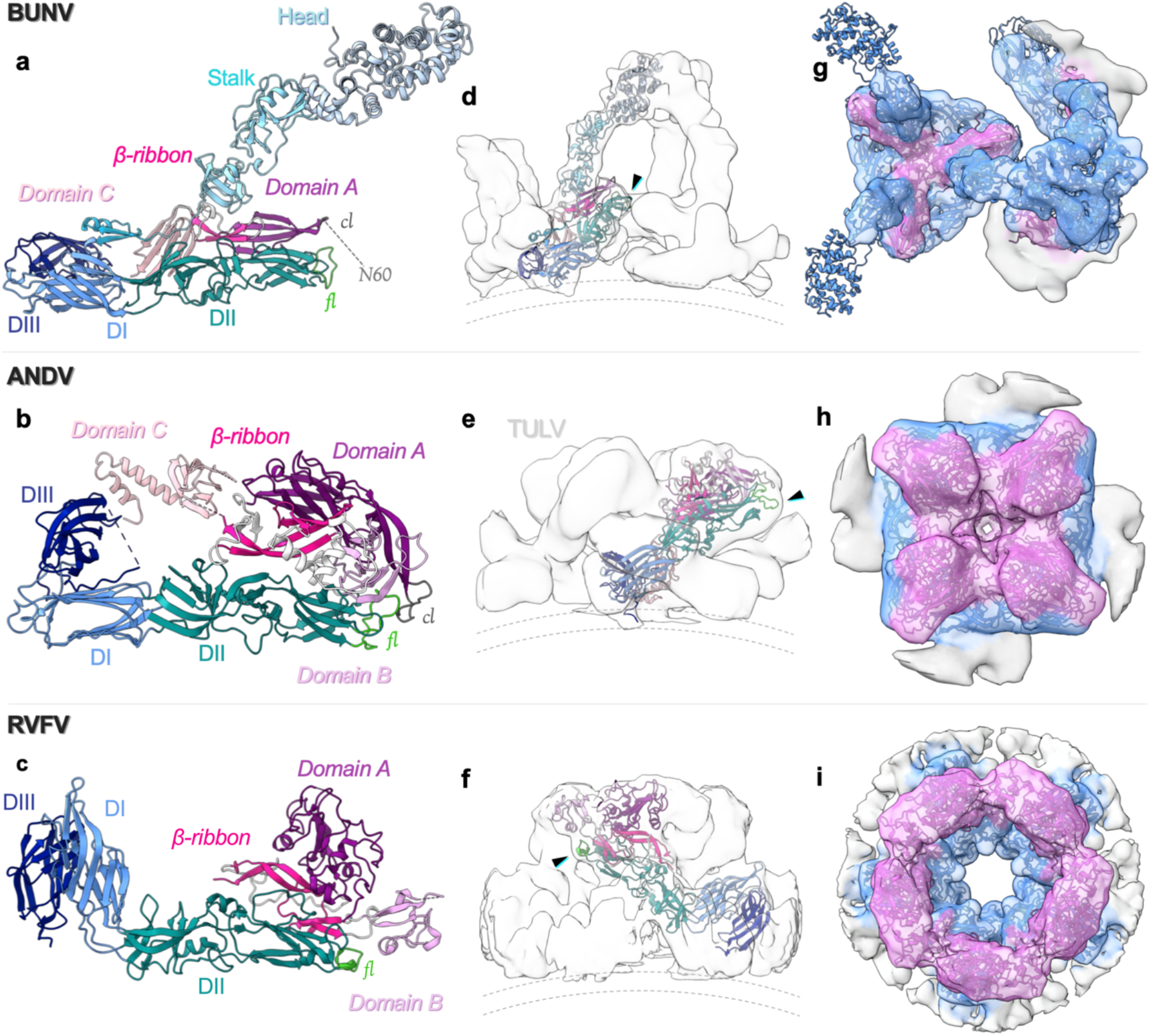
Comparison of GP arrangements of BUNV with that of hantaviruses and phenuiviruses. **a** The BUNV (orthobunyavirus) AlphaFold structure (as in *Fig. 5*) of Gc (shades of blue) and Gn (shades of pink), as compared with the X-ray crystal structures of **b** the hantavirus ANDV (pdb: 6zjm) and **c** the phenuivirus RVFV (in the conformation as fitted within the EM average; pdb: 6f9f). The individual domain colours are correlated between the three models, corresponding to the sub-domains of Gc (head, stalk, domain I (DI), DII, DIII and the fusion loop (*fl*)) and Gn (Domain A, Domain B, Domain C, β-ribbon and the capping loop (*cl*)); as previously described (23,28). The *fl* is shown in green and the *cl* in dark grey for each. **d,e,f** Fit of a single Gn-Gc dimer within the corresponding EM density average; side view. The black arrows indicate the position of the fusion loop and dotted lines the viral membrane. **d** Shows the BUNV AlphaFold fit within the floor and tripod STA (as in *Fig. 5*). **e** ANDV Gn-Gc dimer within TULV STA reconstruction (emd-11236) (26). **f** RVFV Gn-Gc dimer within the EM single-particle reconstruction (emd-4201) (32). **g,h,i** Top-down view of the EM density averages with fitting of the Gn-Gc multimers: **g** BUNV trimer (within the floor and the tripod), **h** ANDV tetramer, **i** RVFV pentamer. The density for each has been coloured by distance from residues of Gn (pink) or Gc (blue), demonstrating the relative positions of each within the complex.

At the time the prediction was made, AlphaFold had not yet been trained on the recently published post-fusion structures of the orthobunyavirus Gc fusion ectodomains (pdb:7A57), a circumstance that allowed further validation of the model. Indeed, the individual Gc fusion domains I, II, and III of the model are virtually identical to those of the LACV x-ray structure with RMSD values of 1.08 Å, 1.16Å, and 0.95 Å (pruned atom pairs), respectively, and in the model are arranged in the canonical pre-fusion orientations relative to one another. The arrangement of the Gc fusion domain in the floor STA shows that a trimer fits well within this average (Fig. 5e,f), and in a highly similar way to that predicted using the LACV structure with the fusion loops pointing upward underneath the tripod (Fig. 4e,f and Supplementary Movie 1).

The Gn AlphaFold model also displays the typical architecture of its orthologs from hantaviruses, phleboviruses, tospoviruses and alphaviruses, which are divided into domain A, β-ribbon, domain B and domain C (28). Similar to Gn from other bunyavirus families, BUNV Gn is formed of β-sheets, corresponding to domains A, C and β-ribbon (28), and sits parallel to domain II of the Gc fusion domain, likely forming numerous stabilising interactions (Fig. 6a). However, the orthobunyavirus Gn is significantly shorter than the orthologues and the model suggests that it lacks the canonical domain B, and instead resides below an additional region from Gc, the stalk (Fig. 6, compare a, b & c). Our unambiguous fit of three Gn-Gc heterodimer models into the triangular floor of the spike also demonstrates that three protomers of Gn trimerize at the centre of the floor in a similar fashion as in the tetrameric hantavirus spike (26) (Fig 6).

Overall, the AlphaFold prediction confirms the orientation and positions of both Gn and Gc, and allows us to model the whole orthobunyavirus GP envelope. This arrangement emphasises the role of Gn in stabilising the tripod and the fusion domains in the floor region, and demonstrates how the lattice-like local arrangements are formed (Fig. 5g,h and Supplementary Movie 2).

### pH 6.3/K^+^ affects the floor region, resulting in GP trimer uncoupling

As we had established an effect of pH 6.3/K^+^ on the floor domain of the BUNV GP, we subsequently used STA to determine the GP arrangement from pH 6.3/K^+^ virions; resolved to ~16 Å resolution (FSC cut-off of 0.5, Supplementary Fig. 4d). A dramatic change in GP architecture was observed, with the lack of clear tripods and the flexibility of the spikes evident early during processing (Fig. 2f,i,j), suggesting that pH 6.3/K^+^ treatment had uncoupled the tripods (Fig. 7a-d). A short spike-like projection was aligned extending from the membrane (~10 nm), with a large, flat region of density above the lipid bilayer, which did not form the regular floor domain structure observed in the pH 7.3/no K^+^ condition. The lack of density at the top of the spike was indicative of a highly dynamic unstructured region, exhibiting multiple conformations which cannot be resolved by STA (Fig. 7a,c). However, two features were apparent from these averages: 1) the lack of organised tripods indicates uncoupling of this region, with the ‘spike’ potentially corresponding to an extended Gc fusion domain or stalk domain from the pH 7.3/no K^+^ average (Supplementary Fig. 7); 2) a floor region was present in this average, albeit it did not appear to form the ordered trimeric arrangement at the base of the spike observed for the pH 7.3/no K^+^ (Fig. 7d,e and Supplementary Movie 1). Interestingly a monomer of the Gc fusion domain in its pre-fusion conformation did not fit well within this region, indicating that it is likely located within the ‘spike’ and extended away from the viral membrane (Supplementary Fig. 7).

**Fig. 7.**
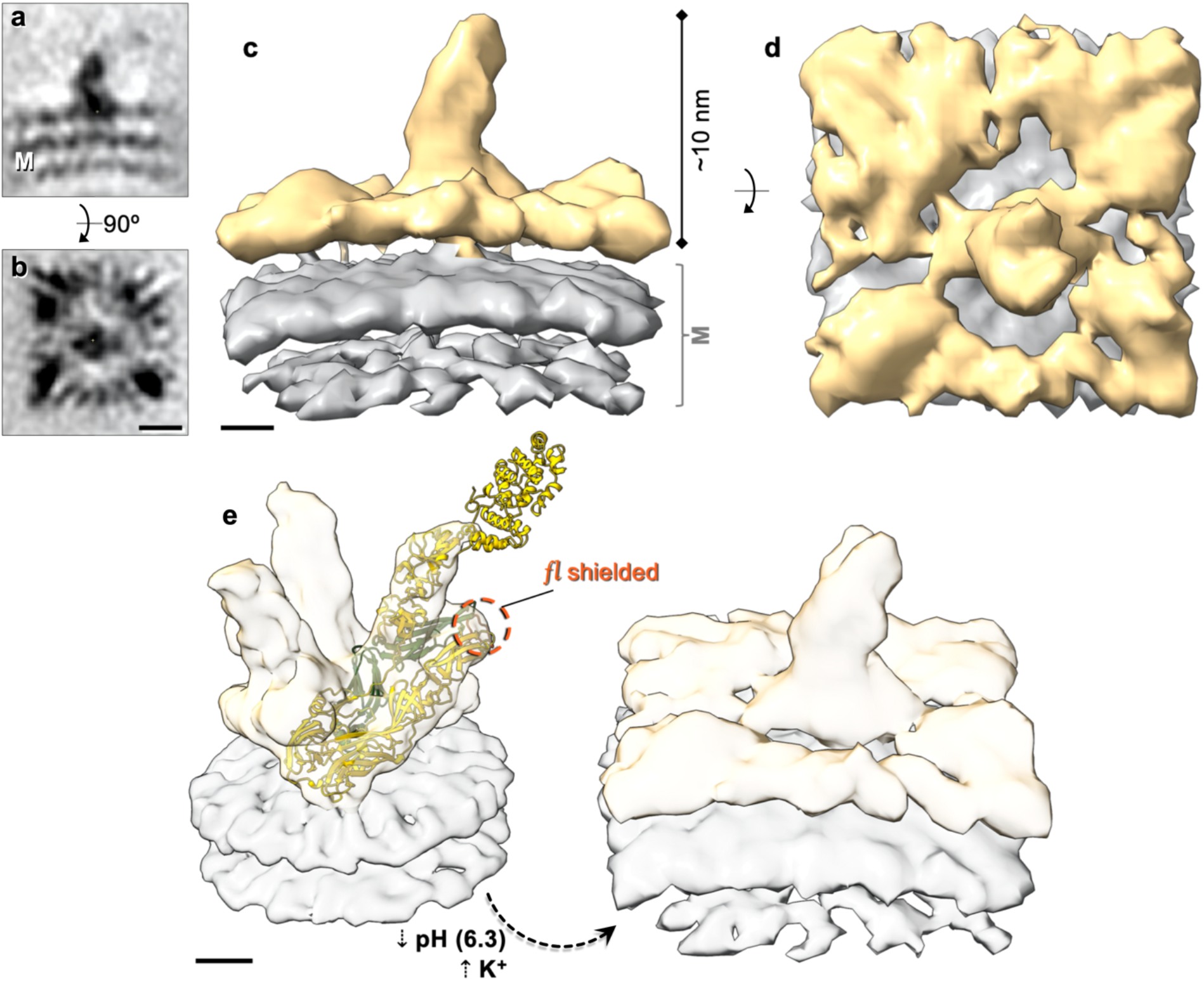
The trimeric architecture of BUNV GPs is abolished by pH 6.3/K^+^ treatment. STA of ~10,000 subtomograms manually selected from the surface of pH 6.3/K^+^ virions (EMD-15579). **a,b** Slices through the centre of the spike electron density from the side (*a*) and top (*b*). The viral membrane is indicated (M). **c,d** Isosurface rendering of the density in *a,b*, with the spike (gold) on top of the viral membrane (grey). **e** Scaled comparison of the pH 7.3/no K^+^ floor region STA with the AlphaFold model fitted, and the pH 6.3/K^+^ STA. The location of the fusion loop (*fl*, orange) is indicated in the pH 7.3/no K^+^ floor STA. Scale bars: *a*, *b* = 5 nm; *c*, *d*, *e* = 2 nm.

With the lack of observable tripods, we took advantage of a monoclonal anti-Gc neutralising antibody (mAb-742) which recognises a conformational epitope on the head domain (near glycosylation site N624, which resides on the top of the head (Fig. 3c,g, N624 indicated)) (29,30). Western blot for BUNV-N of lysates from cells infected with viruses treated at pH 7.3/no K^+^, pH 6.3/no K^+^ or pH 6.3/K^+^, and incubated with mAb-742, showed that this antibody neutralises infection by all treated virions (Supplementary Fig. 8). This confirms that the head-stalk domains are intact and remain attached after pH 6.3/K^+^ treatment; which was unclear from the STA. It is likely that the pH 6.3/K^+^ viruses are still neutralised owing to steric hindrance.

## Discussion

We previously demonstrated that for BUNV low pH and K^+^ treatment, and for HAZV neutral pH and K^+^ treatment, expedites infection. However, mechanistic insight using HAZV was limited owing to the low resolution of the STA and lack of GP structures available (10,13) and completely missing for BUNV. Here, we utilised BUNV to investigate the mechanistic and structural consequences of endosomal pH and K^+^ during bunyavirus entry using cryo-ET and STA.

Using BUNV, orthobunyavirus GPs were previously shown to form a trimeric tripod structure that extends above the membrane-bound floor region, where interactions with neighbouring floor regions generate locally ordered lattice-like arrangements (19). This contrasts the GP organisation for species from other bunyavirus families, which is tetrameric (13,26,31), or icosahedral (32–37).

Here we have improved the average of the BUNV trimer, allowing us to fit structures of the BUNV head and SBV stalk, confirming previous predictions that the fusion domain resides in the floor region (Figs. 3,4) (16,19). Our improved resolution of the floor region, coupled with an AlphaFold model of the complete BUNV Gn-Gc dimer, allowed us to model the whole viral envelope and demonstrates the hetero-hexameric 3x(Gn/Gc) assembly (Fig. 5g,h). Additionally, the fitting of Gc in our subtomogram average shows by exclusion that Gn resides at the centre of the floor domain, likely stabilising this region. This assembly resembles the organization of the Alphavirus envelope, in which three protomers of the class II fusion protein E1 laterally surround a trimeric core of its chaperone E2 (38). A similar assembly, with however different stoichiometry, is also found in the hetero-octameric 4x(Gn-Gc) complex of the orthohantavirus envelope (Fig. 6h) (26). In the pre-fusion structures of related fusion machineries (*Phenuiviridae* (RVFV) and *Hantaviridae* (TULV/ANDV)) (26,32), Gn typically overtops Gc and is thus likely involved in shielding the fusion loops and mediating initial contact with the target cells (23,39) (Fig. 6g-i). In contrast however, our model of the BUNV assembly illustrates that orthobunyavirus Gn cannot completely cover the Gc fusion domain. The capping loop found at the tip of Gn is also shorter than that of hantaviruses, however it still partially shields the fusion loops (Fig. 6). In addition to its polypeptide chain, a strictly conserved N-linked glycan on the capping loop (residue N60) is likely maintained to directly interact with the fusion loops on Gc (26) (as indicated in Figs. 3c and 6a). This Gn glycan was previously shown to be essential for correct GP folding and successful virus rescue, suggesting an important function in the regulation of the pre/post-fusion switch (30). An intriguing difference to these orthologous systems is the relatively small size of the orthobunyavirus Gn (~40-60% smaller, ~30 kDa), with Gc also being ~40-50% larger (~110 kDa) (23,40). The unique N-terminal half of larger orthobunyavirus Gc appears to compensate for the small size of its Gn, being well positioned for receptor scavenging and indirectly shielding the fusion machinery (Figs. 5,6 and Supplementary Movie 1). These head and stalk domains of BUNV Gc are dispensable for infection and are not involved in cell fusion or Golgi trafficking, indicating no role in subsequent stages of infection (20,21). Furthermore, our results show that the head and stalk domains are not required for virus triggering by pH 6.3/K^+^, indicating a specific effect of this condition on the floor region (Fig. 1d-f). Despite its smaller size, orthobunyavirus Gn likely stabilises the centre of the floor region to maintain the arrangement of the Gc fusion domains, similar to that seen for other bunyavirus glycoproteins.

With the Gc tripod and the Gn capping loop including its conserved glycan shielding the fusion loops, disassembly of the tripod is required during fusion, similar to the glycoprotein rearrangement required for exposure of the fusion loops of RVFV, HTNV and ANDV (26,32,41,42). This disassembly is likely caused by a biochemical trigger during entry which would initiate the structural changes in the GPs to expose the fusion loops, permitting their interaction with endosomal membranes. This is what we observed upon pH 6.3/K^+^ treatment, where the organised trimeric arrangement is disassembled, and could involve exposure of the fusion loops to target membranes. This is consistent with the K^+^ effects previously shown for HAZV, where K^+^ treatment triggered large structural changes in the GPs at neutral pH, causing elongation of the GP spikes and their association with co-purified membranes (13). This however differs from previous studies on the BUNV GP trimer that suggested the occurrence of uncoupling at pH 5.1 in the absence of K^+^ (19), since shown to render BUNV non-infectious (10), therefore likely representing a post-fusion conformation. As the requirement of K^+^ for other bunyaviruses has not been explored, the pH 6.3/K^+^ induced loss of the hetero-hexameric arrangement observed for BUNV may differ to the loss of the ordered tetrameric arrangement of the TULV (hantavirus) GPs at pH 5.0 (42), or the elongation and membrane interaction of GP spikes of UUKV (phenuivirus) at the same pH (33).

We predict that at pH 7.3 (no K^+^) the fusion loops are shielded by the tripodal lattice of Gn and Gc (Figs. 3,4). During entry, BUNV reaches an endosomal environment of pH 6.3 and high [K^+^] (Fig. 8), this acts as a partial-trigger initiating changes in the floor region and uncoupling of the tripod (Fig. 7), generating an intermediate disordered state. From the whole virion perspective, this pH 6.3/K^+^ triggering disrupts the locally ordered lattice, allowing free movement of the GPs on the BUNV envelope and polarisation and clustering of the GPs to permit fusion, at which point areas of the membrane become devoid of spikes (Fig. 2f). This state is capable of interacting specifically with cell membranes (Fig. 1b,c) (but not viral membranes, as pH 6.3/K^+^ virions do not interact with each other; Fig. 2d). Similar structural changes in BUNV may also be elicited at low pH alone (pH 6.3/no K^+^), which allows interactions with other virions (Fig. 2c). However, the lack of virus-virus interactions in the presence of K^+^ at pH 6.3 suggests that K^+^ limits the effects of pH and prevents full exposure of the fusion domains. Interestingly, a post-fusion X-ray structure of ANDV Gc at pH 6.5 in the presence of KCl coordinates a K^+^ ion within domain II, at glutamate residue E106, which then forms different interactions and a more unstable arrangement (31). Similar may be true for the pH 6.3/K^+^ BUNV Gc, in which a K^+^ ion coordination may affect exposure of the fusion loops. The fact that K^+^ prevents virion aggregation, but allows interactions with host membranes (Fig. 1b,c), may represent a requirement for a specific host factor. Similarly for HAZV, K^+^ alone was sufficient to induce interactions with membranes, but no fusion events were observed further suggesting the requirement of an unknown host factor (13). Bunyaviruses commonly require additional host factors for fusion events, such as UUKV which requires the presence of a late endosome-resident phospholipid bis(monoacylglycerol)-phosphate, and LACV and DABV which require serine protease activity (33,43).

**Fig. 8.**
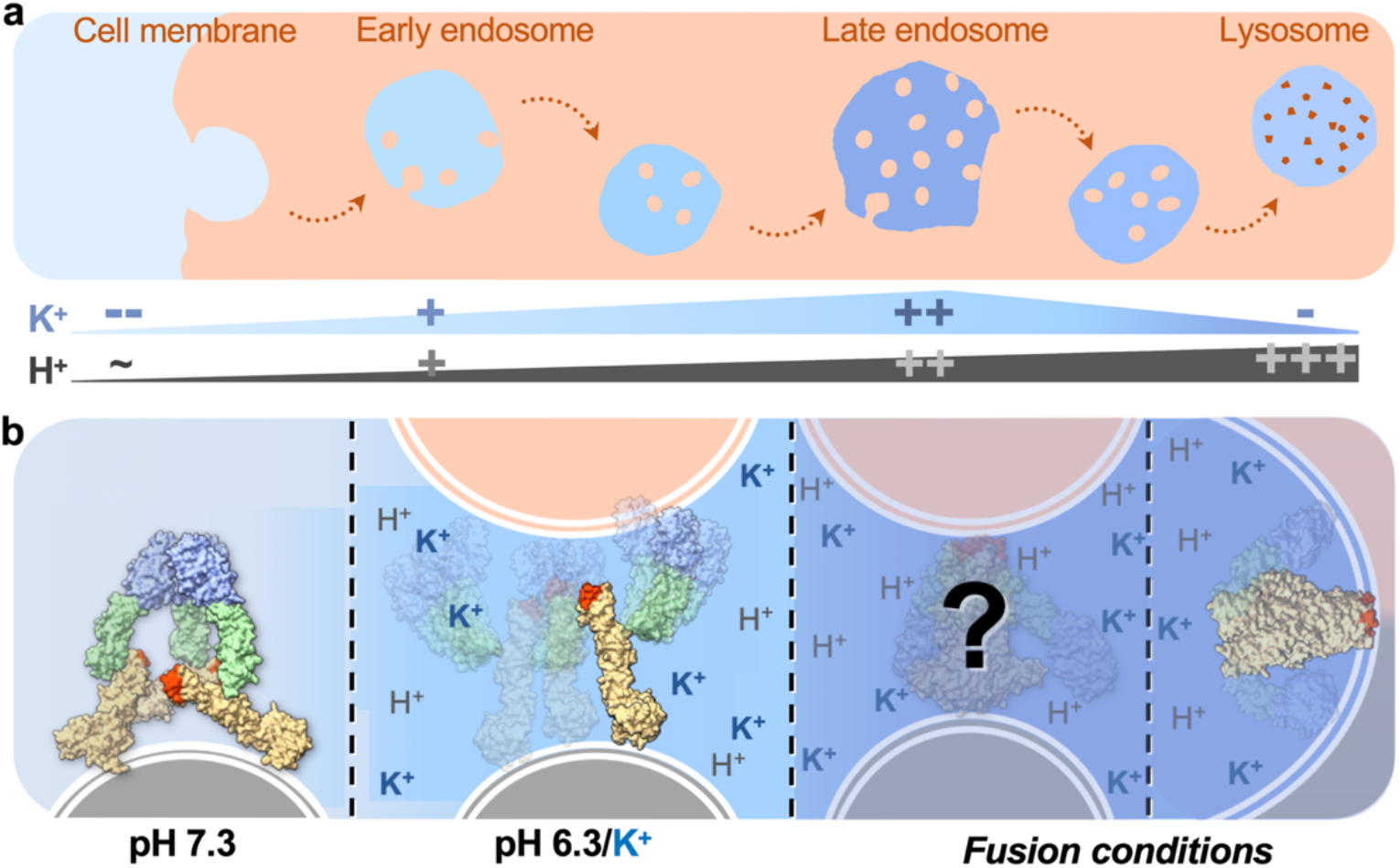
Proposed stages in BUNV fusion during endocytic entry. **a** Model of endocytic transit; from the cell membrane, through early and late endosomes, and into lysosomes. Canonically pH decreases from pH ~7 to pH ~4, whereas [K^+^] is low in the extracellular media and increases with passage into late endosomes, after which decreases again within lysosomes (10,54). **b** BUNV GPs are tripodal and interconnected when entering the cell. During transit, the reducing pH and increasing [K^+^] environment acts as a partial trigger, allowing the tripods to uncouple and exposing the fusion loops. Insertion of fusion loops into host membranes alongside an unknown host factor, and potentially a further reduction in pH, cues subsequent intermediate stages yet to be characterised. This is followed by the canonical fold-back of the fusion domains required to pull the viral and host membranes together (23), resulting in the post-fusion conformation similar to that observed for LACV (pdb:7A57).

In order to transit into a post-fusion conformation, structural changes may also be required in the head and stalk domains. Given that the mAb-742 neutralising antibody, which targets a conformational epitope on the head domain (30), is still effective against pH 6.3/K^+^ treated viruses, we propose that the structure of the head domain is not altered during fusion (Supplementary Fig. 8a-c). In this case, the antibody might work by sterically impeding the refolding of BUNV GPs or blocking the fusion loops from physically reaching the target membrane. Further rearrangements of the fusion domain would then be necessary for the subsequent canonical fusion stages involving viral and host membranes being pulled into close contact, followed by fusion (Fig. 8) (23).

In summary, we have determined the molecular arrangement of the Gc fusion domain and Gn within the floor domain of the BUNV spike, and additionally resolved drastic structural changes in the GP architecture in response to pH 6.3/K^+^, identifying an intermediate fusion state capable of interacting with target membranes. The ability of pharmacological K^+^ channel inhibitors to disrupt infection by a number of bunyaviruses from different families (11,13,14,44), alongside the structural changes elicited by K^+^ on both BUNV and HAZV, suggests a shared mechanism that could be exploited therapeutically.

## Methods

### Cells and viruses

A549 (human alveolar carcinoma epithelial), SW13 (human adrenal cortex carcinoma) and BHK-21 (baby hamster kidney) cells were obtained from the European Collection of Cell Cultures (ECACC). Cells were cultured in Dulbecco’s modified Eagle’s medium (DMEM, Sigma) supplemented with 10% foetal bovine serum (FBS), 100 U/ml penicillin and 100 μg/ml streptomycin, and maintained in a humidified incubator at 37 °C, with 5% CO_2_. Wild-type BUNV stocks (provided by Professor Richard Elliott, Glasgow Centre for Virus Research) were generated from infected cell supernatants and titre estimated by plaque assay as previously described (10,11). GFP-BUNV stocks were provided by Dr Xiaohong Shi (Glasgow Centre for Virus Research) and encode an eGFP tag in place of residues 478-827 of the BUNV M segment (the Gc head and stalk regions) (20). Wild-type human respiratory syncytial virus A2 strain (HRSV) was obtained from the National Collection of Pathogenic Viruses (NCPV) of Public Health England (PHE) and cultured as previously described (45).

### Virus purification

BHK-21 cells were seeded into T175 flasks and infected with BUNV for 3 hrs in FBS-free DMEM (MOI of 0.1). Media was replaced with 2% FBS DMEM and incubated until 44 hpi at 32 °C. Virus-containing supernatants were clarified by centrifugation and filtration (0.22 μm filter), and then virions were pelleted by ultracentrifugation (150,000 x *g*, 3 hrs, 4 °C) through a 30% sucrose cushion (46). Virions were resuspended in 0.1x phosphate buffered saline (PBS, 0.1x to dilute the salts) by gentle rocking at 4 °C overnight. BUNV titres were determined by plaque assay, and purity was determined by silver staining and negative stain EM.

### Plaque assays

Plaque assays were performed in SW13 cells as previously described (10). Viral stocks were serially diluted and SW13 cells were infected and overlayed with semi-solid 1.6% carboxy-methyl cellulose. After 6 days the overlay was removed, cells stained with 1% crystal violet and the number of plaques counted to estimate viral titre.

### Silver staining

During the BUNV purification 10 μl samples were collected of the pre-ultracentrifugation viral supernatant (snt), the post-ultracentrifugation snt, snt-sucrose interface and sucrose. Samples were resolved by SDS-PAGE alongside 0.5 μl of purified BUNV and 1:100 diluted protein ladder, then fixed and stained following the silver staining kit (GE Healthcare) manufacturer’s instructions.

### Western blot

At the indicated time points cells were lysed (25 mM glycerol phosphate, 20 mM tris, 150 mM NaCl, 1 mM EDTA, 1% triton X-100, 10% glycerol, 50 mM NaF, 5 mM Na_4_O_7_P_2_, pH 7.4), and western blotting was performed as previous (10,45). Proteins were resolved by SDS-PAGE and transferred onto polyvinylidene difluoride (PVDF; Millipore) membranes. After blocking in 10% (w/v) milk in TBS-T, the appropriate primary antibodies (sheep α-BUNV-N (1:5000; produced by J N Barr (11)), goat α-HRSV (1:1000; Abcam ab20745), mouse α-GAPDH (1:1000, Santa Cruz sc47724) or mouse α-GFP (1:1000; Santa Cruz sc-9996)) were added at 4 °C overnight, followed by the corresponding HRP-conjugated secondary antibodies (Merck α-sheep A3415; α-goat A8919; α-mouse A4416) for 1 hr. Labelling was detected using enhanced chemiluminescence (Advansta) and a Xograph processor. Densitometry was performed using ImageJ and statistical significance determined using a one-way ANOVA where P<0.05.

### *In vitro* priming assays

BUNV ‘priming’ assays were performed as previously described (10). BUNV was incubated pre-infection at 37 °C for 2 hrs in a range of buffers: pH 7.3 (20 mM tris), and pH 6.3 (30 mM bis-tris) with or without 140 mM KCl (final concentration 128 mM KCl) (‘K^+^’ vs ‘no-K^+^’). Buffer was diluted out with DMEM prior to infection of A549 cells at a MOI of 0.1 for the indicated times (hpi), after which points cells were lysed and BUNV infection assessed by western blot. For the GFP-BUNV, virus was treated with buffers (primed) as with WT BUNV and infection assessed at 18 hpi.

For the cell binding assays, after the pre-infection treatment (pH 6.3/no K^+^, pH 6.3/K^+^, or a no-buffer WT control (Ctl)), buffers were diluted out with cold DMEM and virions added to cells at 4 °C for 1.5 hrs; to allow binding but prevent internalisation (10). Cells were washed 3x 30 secs with 0.1x PBS or left unwashed, and infection was allowed to proceed until 24 hpi (see Fig. 1b). An 18 hpi control was performed alongside (no wash) and lysed at 18 hpi as an internal control (to confirm the increase in BUNV-N with pH 6.3/K^+^ expected at this timepoint). In the initial wash test assay (Supplementary Fig. 1a), A549 cells were infected at 4 °C with BUNV (MOI = 0.1) and left unwashed or washed 3x 30 secs with 0.1x PBS or 0.1% trypsin. Infection was assessed by western blot of cells lysed at 24 hpi.

For the neutralisation assays, BUNV was treated at pH 7.3/no K^+^ (control), pH 6.3/no K^+^ or pH 6.3/K^+^ as above, then buffer was diluted and virions were neutralised with mAb-742 anti-BUNV-Gc monoclonal antibody (1:10,000; produced in the Professor Richard Elliott lab at the Glasgow Centre for Virus Research (29)) or a dH_2_O control for 1 hr with gentle rocking. A549 cells were then infected at an MOI 0.1 for 18 hrs, at which point cells were fixed for immunofluorescence or lysed for western blot. The fixed cells were permeabilised with methanol/acetone, immunofluorescently stained for BUNV-NP (α-sheep Alexa-fluor-594 secondary antibodies; Thermo Scientific A11016) and images taken using an IncuCyte ZOOM live imaging system as previously described (10,45).

### Acid-bypass assay

To induce fusion at the plasma membrane, and thus bypass endosomal entry (see Supplementary Fig. 1b), a protocol was adapted from the procedure previously outlined by Stauffer *et al*. (2014) (18). BUNV (MOI 0.1) was bound to A549 cells by addition at 4 °C for 1 hr, and unbound virus was then removed with the media. Virions bound to the cell membrane were subjected to acidic buffers, acid-bypass, by adding warm fusion buffers (pH 6.3/no K^+^ or pH 6.3/K^+^) to cells for a 2 min pulse at 37 °C (instead of the 2 hrs used for ‘priming’), alongside a DMEM infection control. The pH 6.3 fusion buffers used were the same as those used for the priming in the above experiments, but additionally containing 0.1x PBS to maintain cell osmolarity. Cells were subsequently washed with cold DMEM and each condition (pH 6.3/no K^+^ or pH 6.3/K^+^) was treated with or without drugs to prevent endocytic entry. 10 mM tris(2-carboxyethyl)phosphine (TCEP; Sigma) was added to cells for the ‘+ drugs’ wells for 5 mins at 37 °C to inactivate non-internalised virions (12), which was then removed and replaced with an acid-bypass ‘stop’ buffer (DMEM containing 50 mM HEPES (pH 7.3) and 20 mM ammonium chloride (NH4Cl); Sigma) to prevent entry by endocytosis (10). As such, only virions that bypass the endocytic network and fuse at the plasma membrane would result in a productive infection (10,12). Cells were incubated at 37 °C for 24 hrs, then lysed and infection assessed by western blot. For non-drugged control wells (‘-drugs’) DMEM was added instead of drugs, under the same conditions, and then cells were incubated until 24 hpi; also included were pH 6.3/no K^+^ and pH 6.3/K^+^ acid-bypass control wells to be lysed at 18 hpi to confirm triggering occurs under these conditions as with priming experiments.

### Electron microscopy

Negative stain EM was used to assess the purification of BUNV prior to cryo-EM (as described previously (12)). Briefly, purified BUNV was loaded onto glow-discharged carbon-coated grids and allowed to stand for 30 secs. Grids were washed three times with dH_2_O and stained with 1% aqueous uranyl acetate for 10 secs. Images were collected on a FEI Tecnai T12 electron microscope at 120 kV, using a Gatan Ultrascan 4000 charge coupled device camera and operated between −1 μm and −5 μm nominal defocus.

For cryo-EM and cryo-ET, 2 μl purified BUNV virions (~6.8 x 10^6^ particles) were diluted in 2 μl (1:1) of a range of priming buffers (pH 7.3/no K^+^, pH 7.3/K^+^, pH 6.3/no K^+^, pH 6.3/K^+^; final concentration 127.5 mM KCl) for 2 hrs at 37 °C matching the priming experiments used previously (10). Virions were then mixed with 2 μl of Protein A conjugated with 10-nm colloidal gold (Aurion) as a fiducial marker for tomogram alignment, and 3 μl of the mixture was immediately loaded onto glow-discharged lacey-carbon EM grids. Grids were blotted for 3 secs and vitrified using a Leica EM GP automatic plunge freezer. Cryo-EM micrographs were collected for each condition on an FEI Titan Krios using a Falcon E3C direct electron detector, operated at 300 KeV and at −0.5 to −3.5 μm defocus.

### Cryo-ET and image processing

Grids for cryo-ET were prepared as described above and tilt series were collected using a FEI Titan Krios using an energy-filtered (20 eV slit) Gatan K2 XP summit direct electron detector in counting mode (300 KeV), and a Volta phase plate. Tomography 4 (FEI) software was used to collect single-axis tilt series from −60° to +60°, at 2° increments. A defocus of - 1 μm was used, at 53,000x magnification, corresponding to a pixel size of 2.72 Å, and an electron dose of ~1.8 e^-^ per image (total dose per tilt series ~108 e^-^/Å^2^). Each tilt-series projections was pre-processed by motion correction using Relion 3.0, and contrast transfer function (CTF) calculated using Relion 3.0 (gctf) and then corrected using CTF phase-flip (47,48). The IMOD package eTOMO was used to align the projections using the gold fiducials and calculate the 3D reconstructions by weighted back projection (49), with a final pixel size of 5.44 Å after binning by a factor of two. For STA, unbinned tomograms (2.72 Å pixel size) were also generated using only the −30° to +30° tilt angles (to improve the signal to noise of the resulting tomograms). Representative images were Gaussian Blur 3D filtered using ImageJ.

### Subtomogram averaging

PEET (IMOD package) was used for STA (50) and Bsoft for basic image processing (51). Briefly, for both conditions ~200 spikes were initially selected and the spikeInit programme used to calculate initial orientations perpendicular to the viral membrane. These initial orientations were employed to generate an initial reference used for subsequent particle selection.

For the pH 7.3/no K^+^ STA, the seedSpikes and SpikeInit programmes were used to automatically estimate all spike positions and orientations on the viral membranes (71 virions used, ~9,000 subtomograms), which were then iteratively refined following PEET guidelines using the previously generated initial reference. Duplicate particles were discounted at each stage. After the initial refinements C3 symmetry was evident, therefore in later stages pseudo-particles representing the 3 possible orientations of each tripod were calculated, thereby tripling the number of subtomograms. Additionally, re-creating unbinned tomograms using −30° to +30° tilt angles, to remove the projections from high tilt angles, improved the resolution achievable in STA. For the whole tripod, this resulted in 20,282 subtomograms with a calculated FSC resolution of ~9.1 Å, using the calcFSC PEET programme and a 0.5 cutoff. The subtomogram average of the floor region was generated as above, but centring the average on this region instead of the tripod (using the modifyMotiveList programme) and then iteratively refining the locations and orientations of the subtomograms. This similarly resulted in 16,096 subtomograms, but an improved resolution of ~6.6 Å (0.5 FSC cutoff) of this region using the full dataset.

For accurate structural determination and resolution calculations GS-FSC was also performed on half split datasets. For GS-FSC resolution estimation, all automatically selected particles were passed through an initial iteration to remove duplicate particles. The remaining particles were split into half dataset and independently aligned and averaged following identical steps to that outlined above. The half datasets were then aligned to bring the two averages into a common position and orientation, following PEET guidelines. The FSC between the two half maps was computed using the calcUnbiasedFSC programme (PEET), the result of which was used to filter the resolution of the full dataset. This resulted in an unbiased FSC calculation (using a 0.143 cutoff) for the tripod was ~16 Å and ~14 Å for the floor STA.

For the pH 6.3/K^+^ treated virus STA, automatic selection of spikes using seedSpikes, did not result in a coherent average. Subtomograms were therefore manually selected (~20,000) from 89 virions and alignment was refined iteratively using the initial reference, obtained as for the control. No symmetry was evident and therefore was not applied. Additionally, the density at the top of the spikes could not be well refined suggesting multiple conformations. To address this, principal components analysis (pca) and clusterPca programmes were used to identify clusters of separate conformations. Three clusters were identified however the top of the spikes remained unresolved for all (this was also the case for all the clustering options explored) and therefore suggested a highly dynamic structure. The largest cluster (9,987 subtomograms) was taken forward for further refinements utilising a loosely fitted mask to improve the more structured regions and resulted in ~16 Å resolution. This was determined using the calcFSC (0.5 cutoff) PEET programme (not GS-FSC), owing to the high flexibility and low resolution of the average.

Subtomograms were visualised using Chimera and ChimeraX (52,53). Fitting of the BUNV head domain (pdb: 6H3V), SBV stalk (pdb: 6H3S) (16) and the LACV fusion domains (pre-fusion modelled from pdb: 7A57) was also performed in Chimera X. Comparison of the RMSD values comparing the individual domain structures of the LACV and AlphaFold Gc fusion domains was obtained using the matchmaker command in Chimera X. RMSD values for the pruned atom pairs and all atom pairs were obtained: DI pruned pairs (65 atoms) 1.08 Å and all pairs (122 atoms) 5.83 Å; DII pruned pairs (137 atoms) 1.16 Å and all pairs (186 atoms) 1.91 Å; DIII pruned pairs (78 atoms) 0.95 Å and all pairs (84 pairs) 1.16 Å.

AlphaFold v2.1 was used to generate the multimer structure of the BUNV Gn/Gc complex, using the M segment polyprotein (Gn-NSm-Gc) sequence for Gn and Gc from ncbi (17,27). At the time of model generation, only the Gc head and stalk domains were available for orthobunyaviruses; no Gc fusion domain nor Gn structures were available for training the model. For fitting into the tripod STA, the output model was rotated about the flexible inter-domain regions (which also have low model confidence) head-stalk and stalk sub-domains I-II (see Supplementary Fig. 6c).

## Data Availability

The STA structures of the (pH 7.3/no K^+^) BUNV tripod (accession number: EMD-15557), floor region (EMD-15569) and pH 6.3/K^+^ GPs (EMD-15579) have been deposited in EMBD. The AlphaFold model, lacking the TMDs (which are not supported by the STA data) can be found as Supplementary Material 1. Correspondence of materials: John N. Barr.

## Acknowledgements

Thanks to the Astbury Biostructure Laboratory electron microscopy facility for support in using the microscopes, and a particular thanks to Dr Emma Hesketh, Dr Rebecca Thompson and Dr Daniel Maskell (University of Leeds) for the cryo-electron tomography data collection setup. Thanks to Professor Félix Rey (Institut Pasteur, Paris) for kindly providing the structure of the LACV post-fusion Gc prior to publication and for useful comments on the manuscript. Thanks also to Dr Xiaohong Shi, Professor Richard Elliott and Professor Alain Kohl from the Glasgow Centre for Virus Research (CVR, University of Glasgow), and Dr Cheryl Walter (University of Hull) for providing reagents.

## Supporting Figure Legends

**Supplementary Fig. 1.**
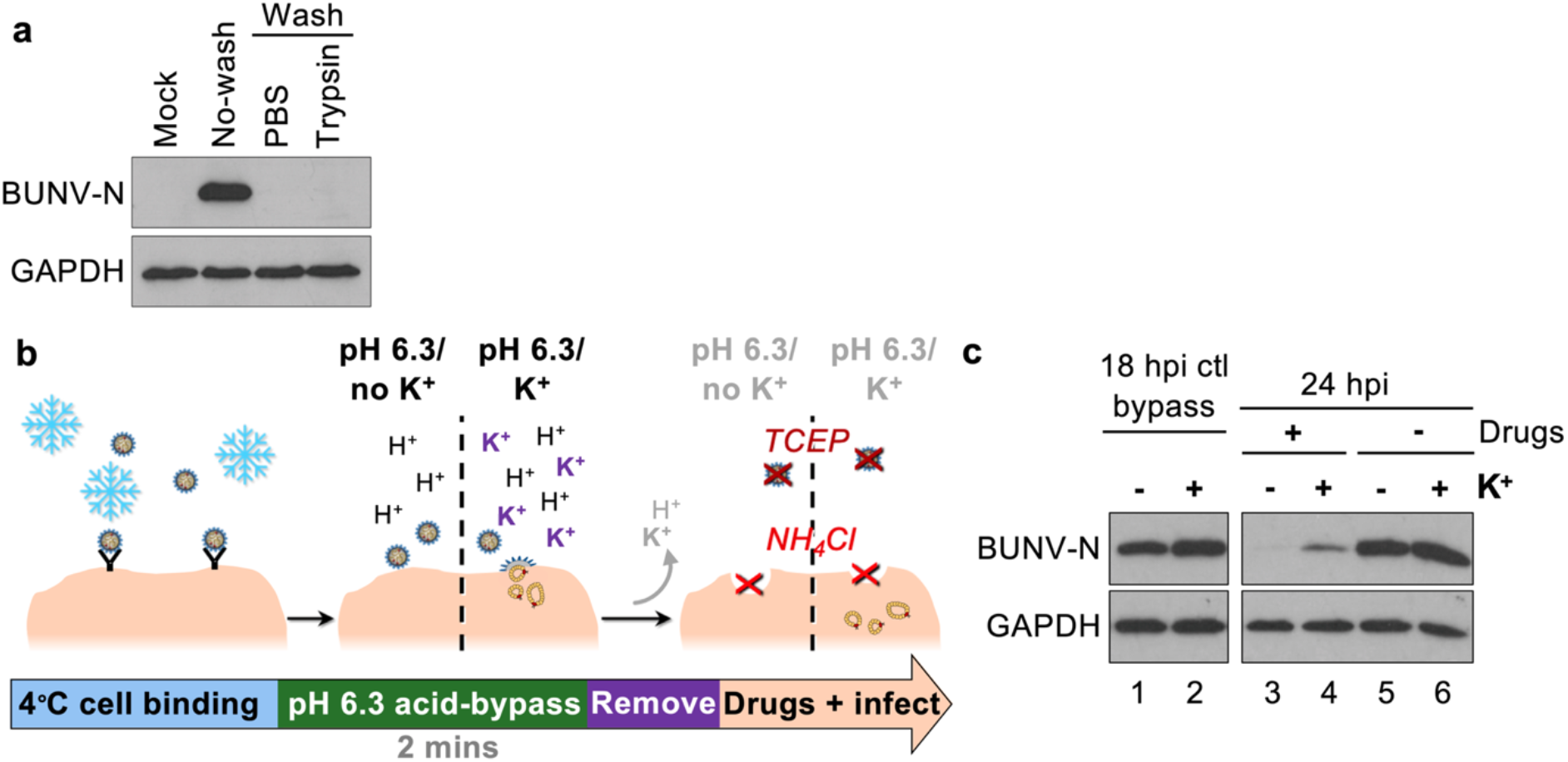
pH 6.3/K^+^ treatment of BUNV increases interactions with host cells. **a** Virions were bound to cells at 4 **°**C and then weakly bound or unbound virions were removed with three PBS or 0.1x trypsin washes (or no-wash as a control). Cells were warmed to 37 °C and infected (or non-infected, mock) for 24 hrs. Western blot analysis of cell lysates (as in *Fig. 1*) using anti-BUNV-N antibodies revealed lack of N signal in the washed samples indicated virions had been removed from the cell membrane. **b** Workflow of an acid-bypass assay. Virions are bound to cells at 4 **°**C and fusion at the plasma membrane is attempted by acid-bypass at pH 6.3/no K^+^ (-) or pH 6.3/K^+^ (+) for 2 mins. Bypass buffer is replaced with media with or without drugs to prevent endocytic entry (TCEP and NH4Cl). Cells are lysed at 18 hpi (internal control) or 24 hpi and BUNV-N assessed by western blot. **c** Western blot of experiment in *b*, where a BUNV-N signal in the presence of drugs (lanes 3-4) indicates conditions where virions were able to fuse with the plasma membrane.

**Supplementary Fig. 2.**
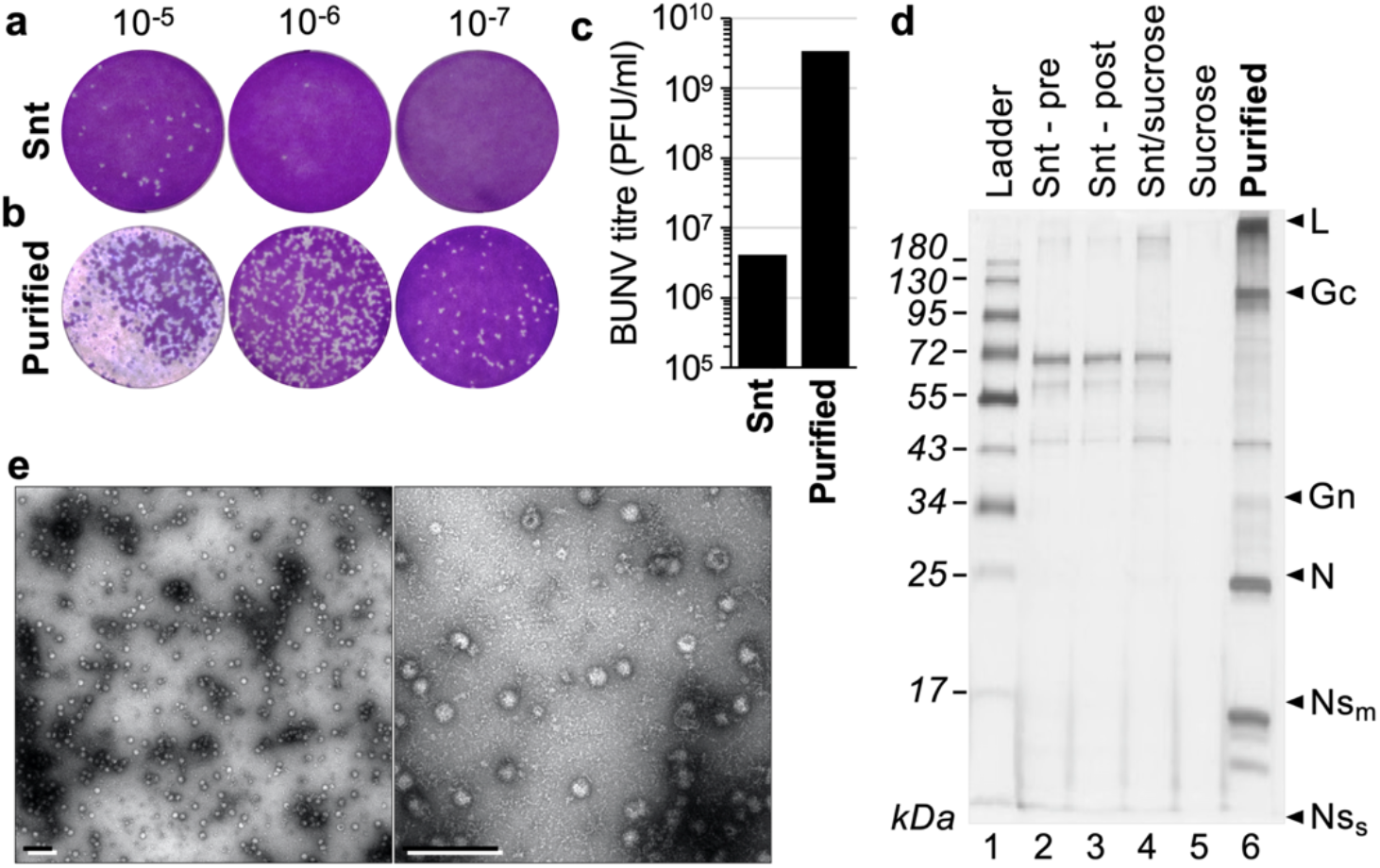
WT BUNV purification. BHK-21 cells were infected with BUNV for 44 hrs and then viral supernatants (snt) were collected and purified by pelleting through a 30% sucrose cushion. Virus was resuspended in 0.1x PBS (Purified) and **a,b,c** the titre was determined by plaque assay as plaque forming units per ml (PFU/ml). **d** SDS-PAGE followed by silver staining of purified BUNV alongside samples collected at each stage of purification; pre-ultracentrifugation supernatant (Snt – pre), post-ultracentrifugation supernatant (Snt – post), supernatant-sucrose cushion interface (Snt/sucrose), and sucrose cushion (Sucrose). The protein ladder sizes are indicated (kDa), as well as the predicted migration of each viral protein based on molecular weight. **e** Negative stain EM of purified BUNV loaded onto carbon coated EM grids and stained with 1% uranyl acetate. Scale bars = 500 nm.

**Supplementary Fig. 3.**
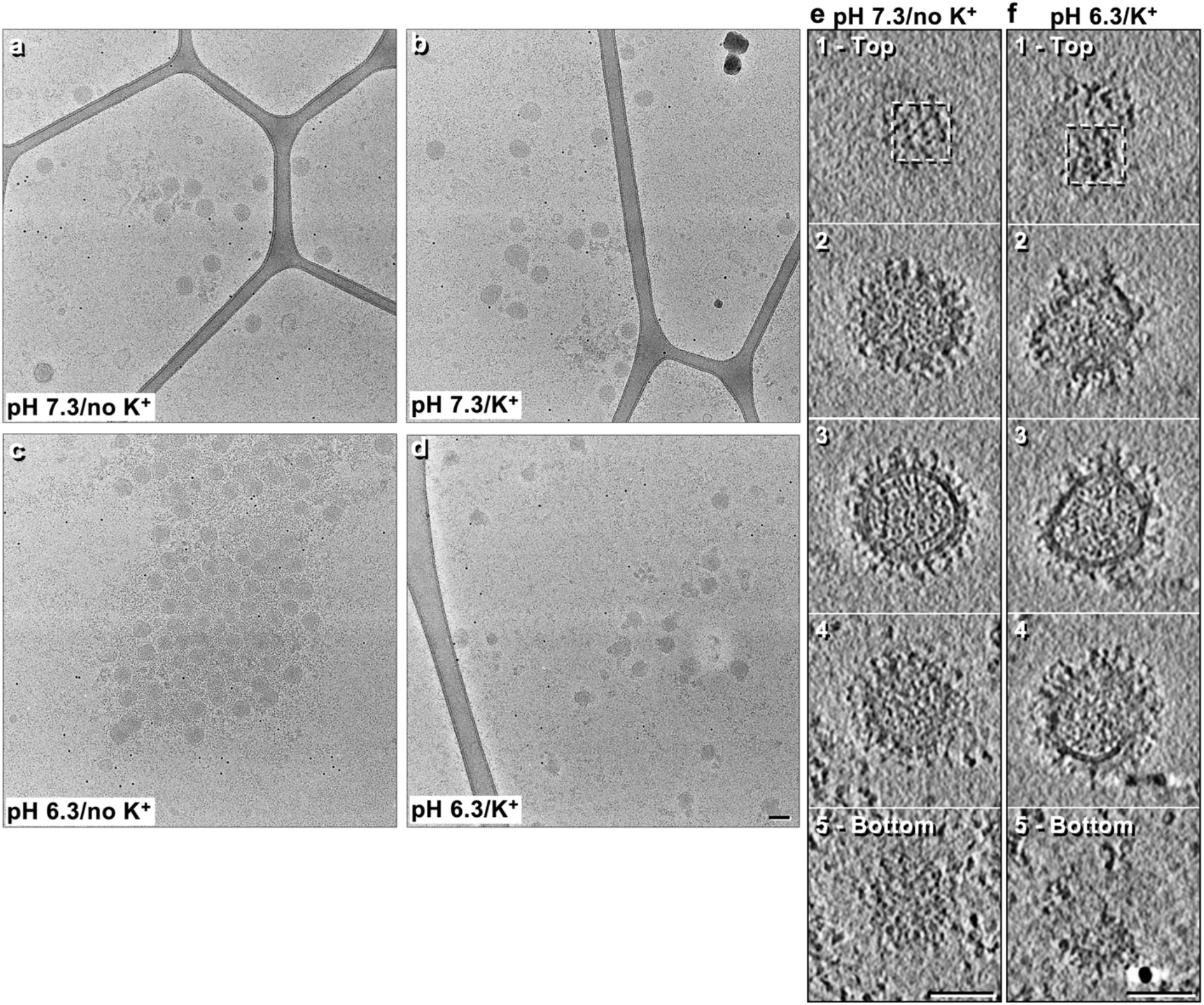
Cryo-EM of BUNV virions under different pH and K^+^ conditions. Widefield micrographs of images shown in *Fig. 2*. **a** pH 7.3/no K^+^, **b** pH 7.3/K^+^, **c** pH 6.3/no K^+^ and **d** pH 6.3/K^+^ treated virions. **e,f** Slices through the z plane of the cryo-ET (as in *Fig. 2e-j*) of individual pH 7.3/no K^+^ (*e*) and pH 6.3/K^+^ (*f*) virions. Scale bars = 100 nm.

**Supplementary Fig. 4.**
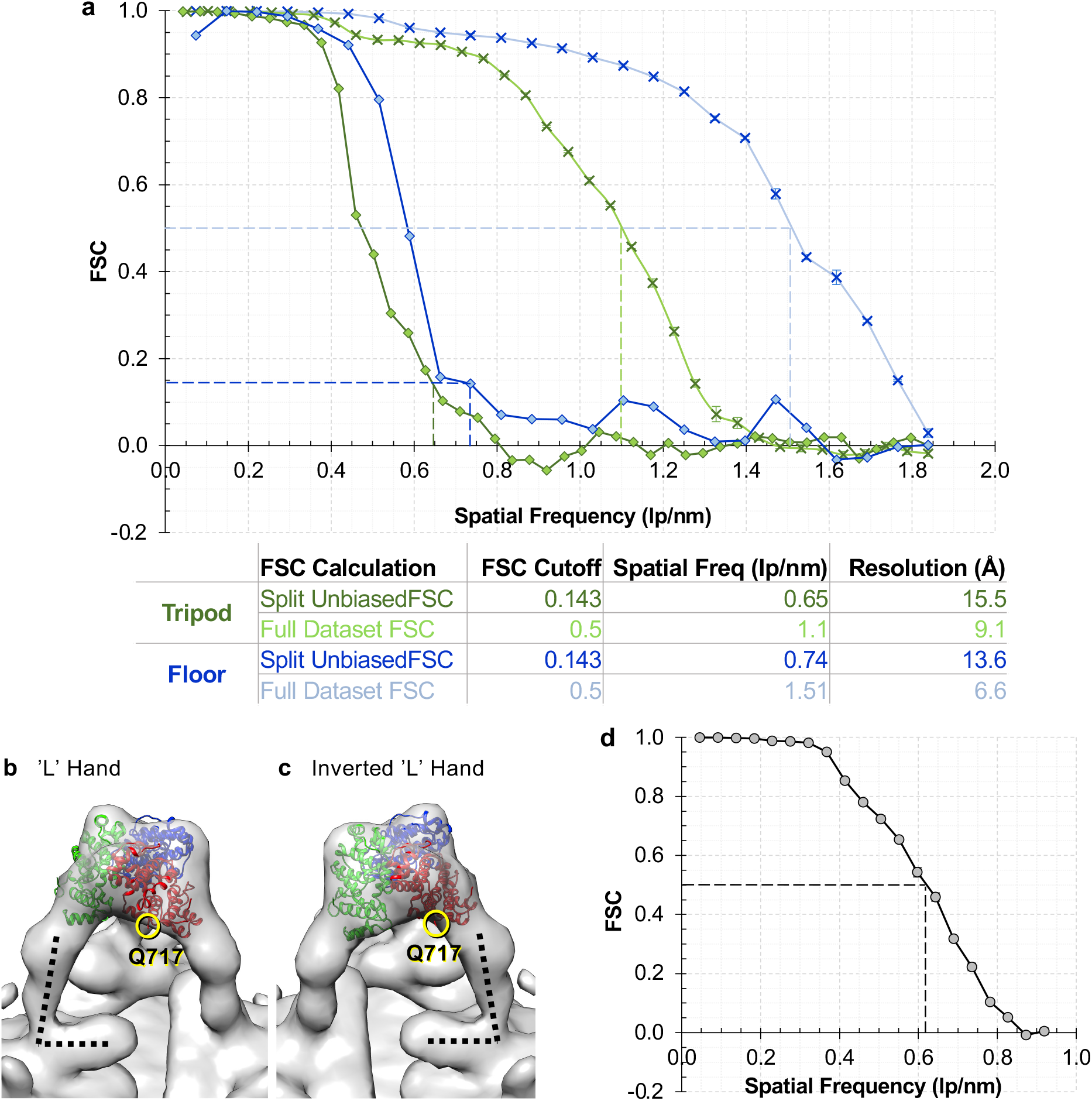
Further analysis of the BUNV GP STA. **a** Fourier shell correlation (FSC) resolution curves of the pH 7.3/no K^+^ BUNV tripod (green) and floor (blue) STAs (*Figs. 3,4*). Gold-standard FSC (GS-FSC) calculations (♦ points) were performed by comparing the resolution of half datasets, using a 0.143 FSC cutoff to determine the resolution of each. Standard FSC calculations (× points) were additionally performed to determine the resolution of full datasets using a 0.5 cutoff. Error bars represent standard deviation from the average of n=3 FSC calculations with different pseudo-random assortment of particles into ‘half sets’ using PEET. **b,c** Fit of the BUNV head domain trimer when switching the handedness; *b* ‘L’ hand, *c* inverted ‘L’ hand. Q717 is the C-terminal residue of the head domain which connects to the stalk domain, and is more favourably localised in the inverted ‘L’ hand (*c*). **d** Standard FSC resolution curve of the pH 6.3/K^+^ BUNV STA. The resolution is ~16 Å (*fig. 7*), using a 0.5 cutoff (used owing to the flexibility of the structure).

**Supplementary Fig. 5.**
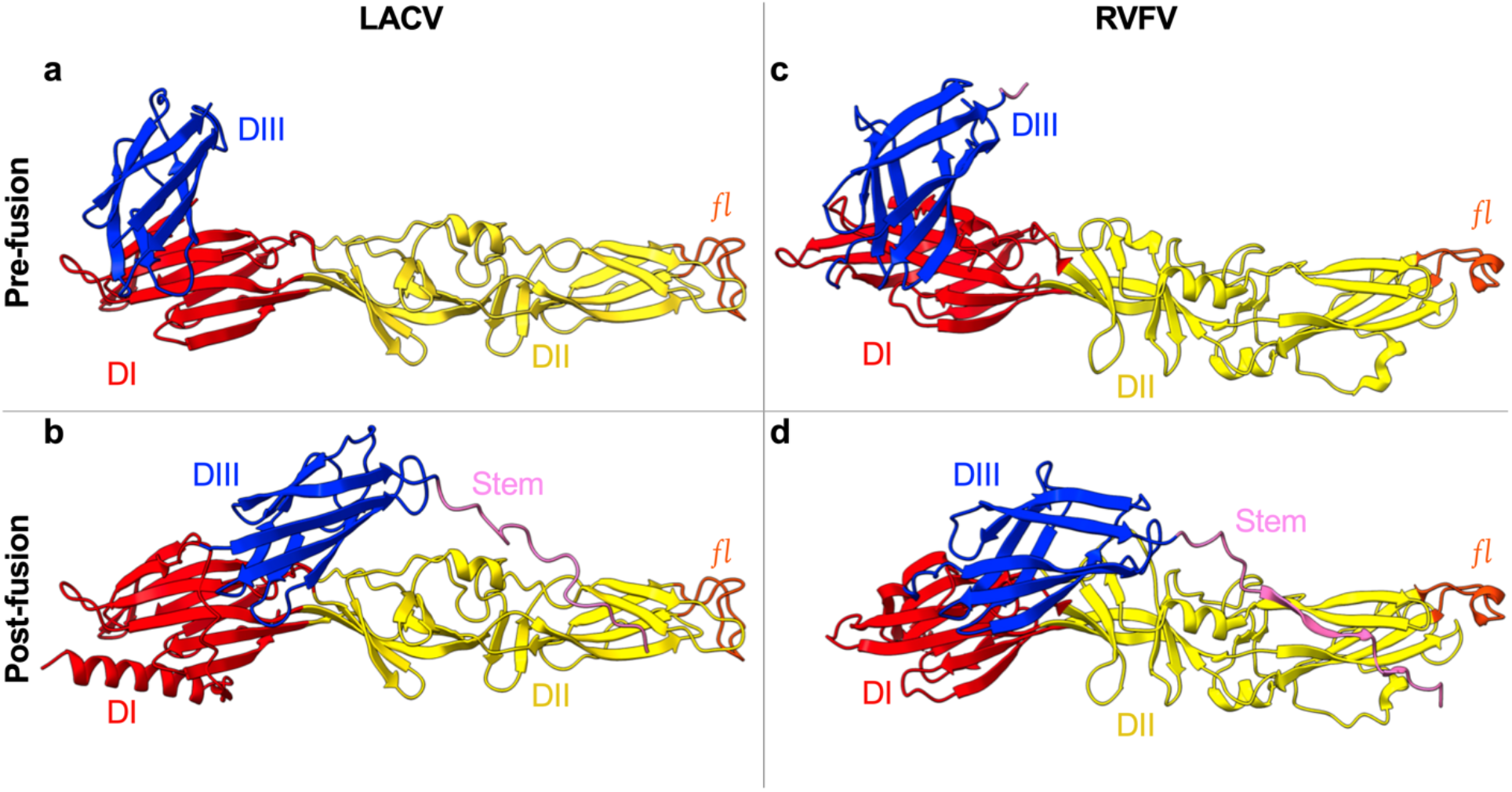
Comparison of the LACV and RVFV pre-fusion versus post-fusion class II fusion domain conformations. A monomer of each conformation is represented, where Domain I (DI) is in red, domain II (DII) yellow, domain III (DIII) in blue, the fusion loop (*fl*) in DII is in orange, and the stem region (connecting to the TMD) is in pink. The **a** LACV pre-fusion fusion domain structure (residues 965-1344) is modelled from the **b** post-fusion LACV fusion domain structure (residues 928-1364; pdb: 7A57). DIII in *a* has been rotated from its position in *b*, based upon the canonical shift in DIII observed for class II fusion proteins in pre-fusion versus post-fusion conformations. RVFV is shown to exemplify the known shift in DIII during fusion events. RVFV fusion domain also forms a class II fold and structures have been solved for both the **c** pre-fusion (residues 688-1118; pdb: 4HJ1 (24)) and **d** post-fusion conformations (residues 691-1136; pdb: 6EGU (25)). The RVFV pre-fusion conformation (*c*) is represented as in the X-ray structure when Gc is expressed recombinantly outside the context of a virus (*Fig. 6c* represents RVFV Gc flexibility fitted within the EM average of the virus (32)).

**Supplementary Fig. 6.**
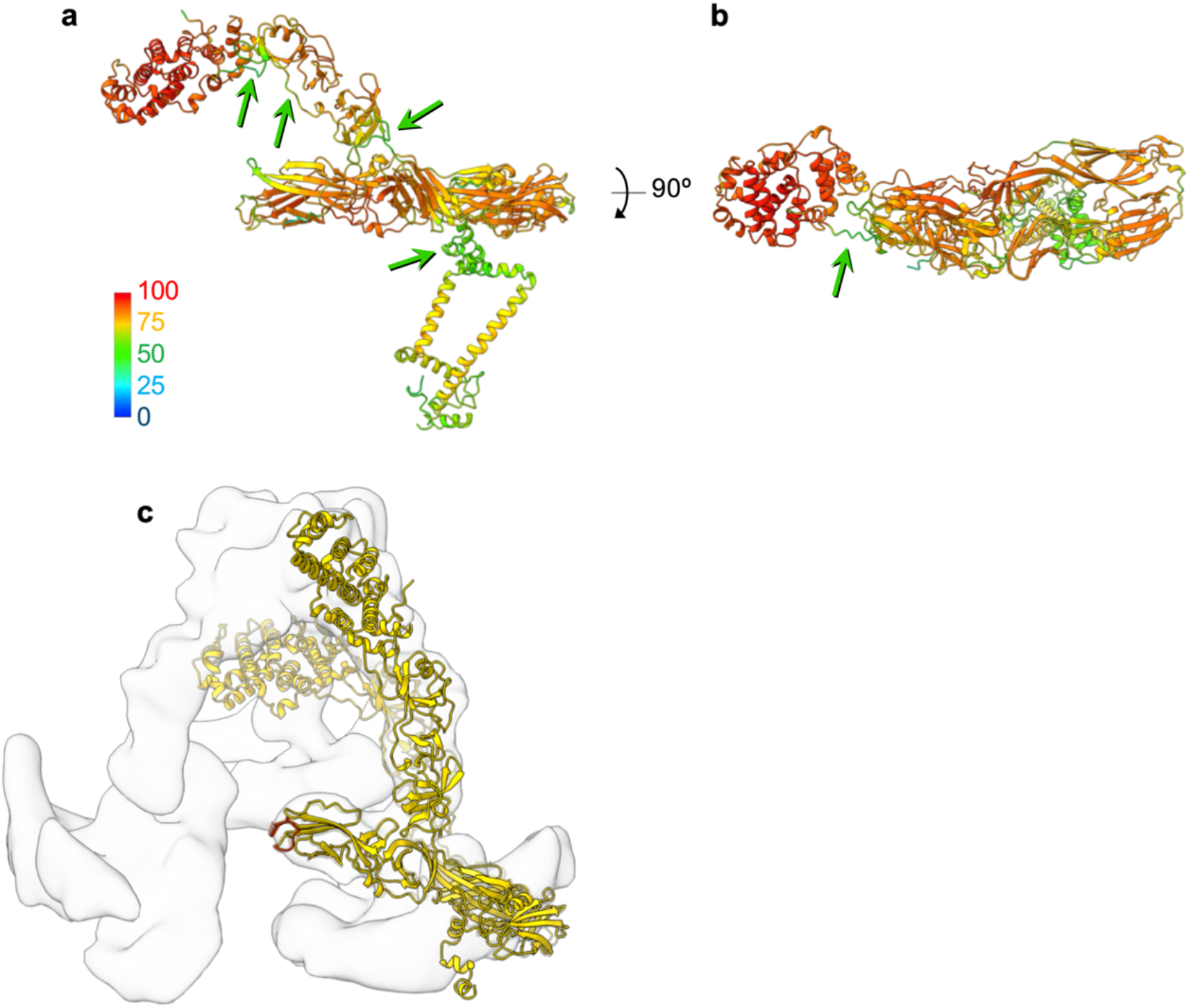
AlphaFold modelling of BUNV Gn-Gc reveals the arrangement of the orthobunyavirus GP envelope. **a,b** The BUNV Gn-Gc model generated by AlphaFold, as in *Fig. 5a,b*, coloured by the per-residue confidence score from 1-100 pLDDT. The majority of residues have a confidence >70 (yellow), indicating the residues were modelled well. Residues >90 are expected to be modelled to high accuracy (17). Key linker-regions have lower confidence (<50) suggesting flexibility between domains compared to the generated model (green arrows). **c** Comparison of the positions of the head and stalk domains in the generated AlphaFold model (translucent), with the fitted model rotated about the head-stalk and stalk sub-domain I-II interfaces.

**Supplementary Fig. 7.**
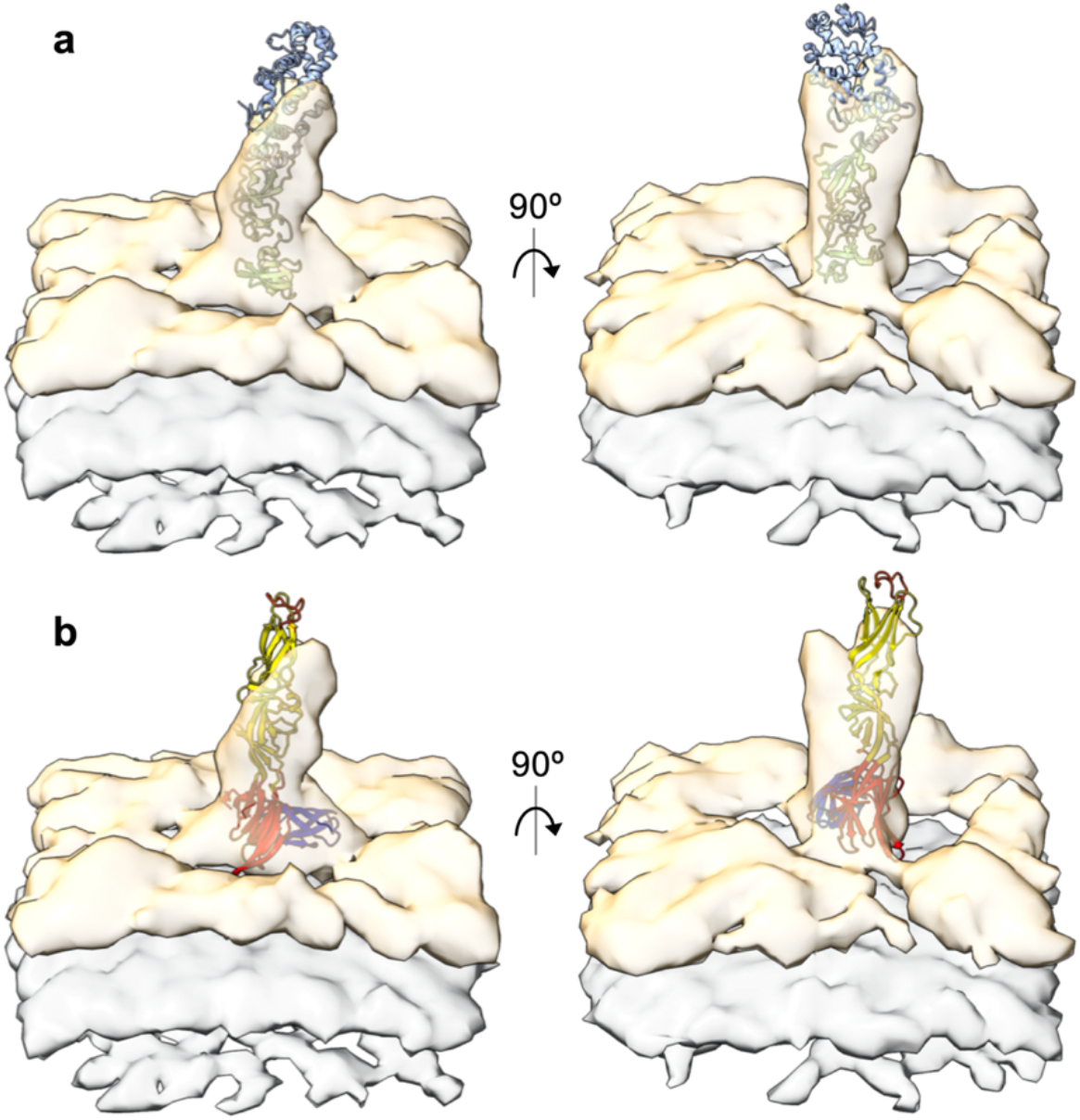
Fitting of solved orthobunyavirus Gc crystal structures on the pH 6.3/K^+^ BUNV STA spike. **a** The BUNV head (light blue, pdb: 6H3V) and SBV stalk (light green, pdb: 6H3S) domains fitted into the spike density. **b** The LACV fusion domain (domains I-III coloured as in *Supplementary Fig. 4*) rotated perpendicular to the viral membrane (grey). DIII (dark blue) remains in proximity to the viral membrane (DIII connects to the C-terminal TMD), with the fusion loops (orange) extending away from the virus.

**Supplementary Fig. 8.**
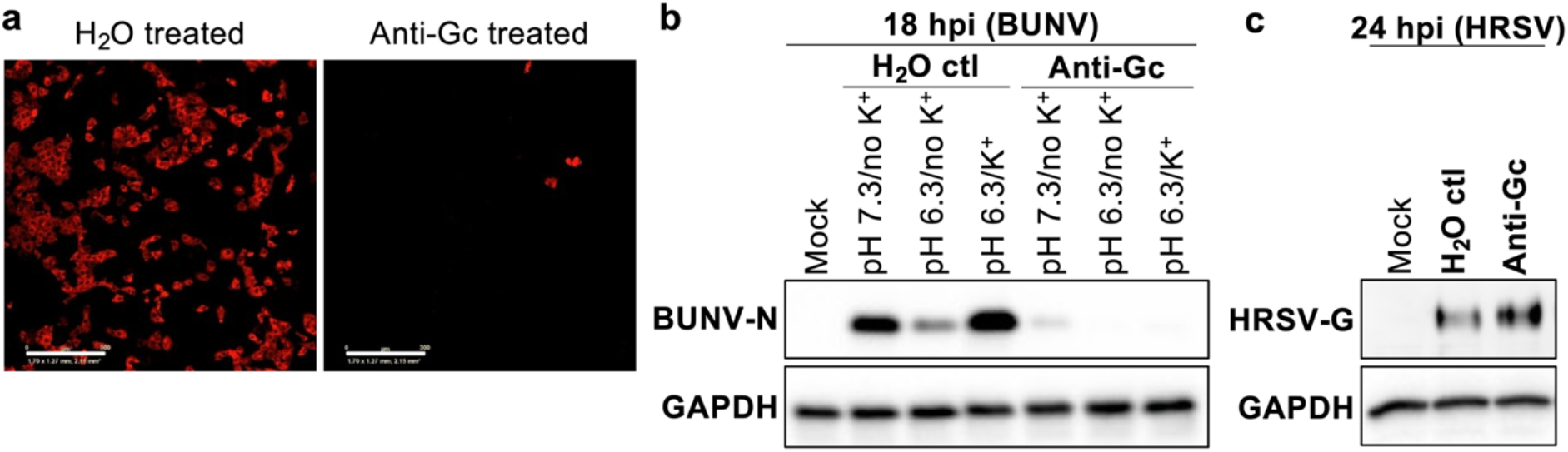
Biochemical characterisation of the Gc head domain after pH 6.3/K^+^ treatment. **a** BUNV virions were treated with dH_2_O (control) or an anti-Gc neutralising antibody that binds the Gc head domain (mAb-742 (29)) for 1 hr. Virus was then added to A549 cells and incubated for 18 hrs, after which cells were fixed, permeabilised and stained for anti BUNV-N with Alexa-Fluor-594 secondary antibodies. Fluorescence images were acquired using an IncuCyte Zoom, identifying fluorescent infected cells. **b** BUNV was pre-infection treated (‘primed’; as in *Fig. 1*), with pH 7.3/no K^+^, pH 6.3/no K^+^ (controls) or pH 6.3/K^+^. Buffer was diluted out and virions neutralised with anti-Gc or a dH_2_O control, and A549 cells infected as in *a*. An uninfected control, mock, was also included. Western blot analysis was performed as in *Fig. 1* using anti-BUNV-N and anti-GAPDH (loading control) antibodies. **c** As a control, HRSV was ‘neutralised’ with anti-Gc or dH_2_O as in *a* and infection was assessed at 24 hpi using an anti-HRSV antibody, to confirm antibody specificity.

## Supporting Movie Legends

**Supplementary Movie 1** Fit of available orthobunyavirus partial-Gc structures and of the BUNV Gn-Gc AlphaFold model within the tripod and floor STA. The BUNV head (6H3V), SBV stalk (6H3S) and LACV fusion domain are fitted as in *Figs. 3,4*; and the BUNV AlphaFold as in *Fig. 5*. CL = capping loop on Gn, FL = fusion loop on Gc. The floor STA is also compared with the pH 6.3/K^+^ virus ‘spike’ as in *Fig 7e*.

**Supplementary Movie 2** Modelling the BUNV envelope. An AlphaFold model of the BUNV GP envelope is predicted, showing three hetero-hexamers fitted within both the tripod and the floor region STA. As in *Fig. 5g,h*. The wider lattice-like model of the viral envelope can be inferred from these arrangements.

## Supporting Material Legend

**Supplementary Material 1** BUNV Gn and Gc ectodomain AlphaFold prediction as is fitted within the BUNV tripod STA; TMDs removed as they do not comply with the STA density (represented are residues Gn 1-187, Gc 1-899).

## References

1. Briese T, Calisher CH, Higgs S. Viruses of the family Bunyaviridae: are all available isolates reassortants? Virology. 2013 Nov;446(1–2):207–16.

2. Elliott RM. Emerging viruses: the Bunyaviridae. Mol Med. 1997;3(9):572.

3. Abudurexiti A, Adkins S, Alioto D, Alkhovsky S V., Avšič-Županc T, Ballinger MJ, et al. Taxonomy of the order Bunyavirales: update 2019. Arch Virol. 2019 May 7;1–17.

4. Zeller H, Bouloy M. Infections by viruses of the families Bunyaviridae and Filoviridae. Rev Sci Tech. 2000;19(1):79–91.

5. Huotari J, Helenius A. Endosome maturation. EMBO J. 2011 Aug 31;30(17):3481.

6. Harrison SC. Viral membrane fusion. Virology. 2015 May 1;479–480:498–507.

7. Halldorsson S, Behrens A-J, Harlos K, Huiskonen JT, Elliott RM, Crispin M, et al. Structure of a phleboviral envelope glycoprotein reveals a consolidated model of membrane fusion. Proc Natl Acad Sci U S A. 2016 Jun 28;113(26):7154–9.

8. Acuña R, Bignon EA, Mancini R, Lozach P-Y, Tischler ND. Acidification triggers Andes hantavirus membrane fusion and rearrangement of Gc into a stable post-fusion homotrimer. J Gen Virol. 2015 Nov 1;96(11):3192–7.

9. de Boer SM, Kortekaas J, Spel L, Rottier PJM, Moormann RJM, Bosch BJ. Acid-Activated Structural Reorganization of the Rift Valley Fever Virus Gc Fusion Protein. J Virol. 2012 Dec 15;86(24):13642–52.

10. Hover S, Foster B, Fontana J, Kohl A, Goldstein SAN, Barr JN, et al. Bunyavirus requirement for endosomal K+ reveals new roles of cellular ion channels during infection. PLOS Pathog. 2018 Jan 19;14(1):e1006845.

11. Hover S, King B, Hall B, Loundras E-A, Taqi H, Daly J, et al. Modulation of potassium channels inhibits bunyavirus infection. J Biol Chem. 2016 Dec 16;291(7):3411–22.

12. Charlton FW, Hover S, Fuller J, Hewson R, Fontana J, Barr JN, et al. Cellular cholesterol abundance regulates potassium accumulation within endosomes and is an important determinant in bunyavirus entry. J Biol Chem. 2019 May 3;294(18):7335–47.

13. Punch EK, Hover S, Blest HT, Fuller J, Hewson R, Fontana J, et al. Potassium is a trigger for conformational change in the fusion spike of an enveloped RNA virus. J Biol Chem. 2018 Apr 20;293(26):9937–44.

14. Sandler ZJ, Firpo MR, Omoba OS, Vu MN, Menachery VD, Mounce BC. Novel Ionophores Active against La Crosse Virus Identified through Rapid Antiviral Screening. Antimicrob Agents Chemother. 2020 Jun 1;64(6).

15. Windhaber S, Xin Q, Uckeley ZM, Koch J, Obr M, Garnier C, et al. The Orthobunyavirus Germiston Enters Host Cells from Late Endosomes. J Virol. 2022 Mar 9;96(5).

16. Hellert J, Aebischer A, Wernike K, Haouz A, Brocchi E, Reiche S, et al. Orthobunyavirus spike architecture and recognition by neutralizing antibodies. Nat Commun. 2019 Dec 20;10(1):879.

17. Jumper J, Evans R, Pritzel A, Green T, Figurnov M, Ronneberger O, et al. Highly accurate protein structure prediction with AlphaFold. Nature. 2021 Jul 15;596(7873):583–9.

18. Stauffer S, Feng Y, Nebioglu F, Heilig R, Picotti P, Helenius A. Stepwise priming by acidic pH and a high K+ concentration is required for efficient uncoating of influenza A virus cores after penetration. J Virol. 2014 Nov;88(22):13029–46.

19. Bowden TA, Bitto D, McLees A, Yeromonahos C, Elliott RM, Huiskonen JT. Orthobunyavirus ultrastructure and the curious tripodal glycoprotein spike. PLOS Pathog. 2013;9(5):e1003374.

20. Shi X, van Mierlo JT, French A, Elliott RM. Visualizing the replication cycle of bunyamwera orthobunyavirus expressing fluorescent protein-tagged Gc glycoprotein. J Virol. 2010;84(17):8460–9.

21. Shi X, Goli J, Clark G, Brauburger K, Elliott RM. Functional analysis of the Bunyamwera orthobunyavirus Gc glycoprotein. J Gen Virol. 2009 Oct;90(Pt 10):2483–92.

22. Hopkins FR, Álvarez-Rodríguez B, Heath GR, Panayi K, Hover S, Edwards TA, et al. The Native Orthobunyavirus Ribonucleoprotein Possesses a Helical Architecture. Brennan B, Cherry S, editors. MBio. 2022 Jun 28;

23. Hulswit RJG, Paesen GC, Bowden TA, Shi X. Recent advances in bunyavirus glycoprotein research: Precursor processing, receptor binding and structure. Vol. 13, Viruses. MDPI AG; 2021.

24. Dessau M, Modis Y. Crystal structure of glycoprotein C from Rift Valley fever virus. Proc Natl Acad Sci U S A. 2013 Jan 29;110(5):1696–701.

25. Guardado-Calvo P, Atkovska K, Jeffers SA, Grau N, Backovic M, Pérez-Vargas J, et al. A glycerophospholipid-specific pocket in the RVFV class II fusion protein drives target membrane insertion. Science (80-). 2017 Nov 3;358(6363):663–7.

26. Serris A, Stass R, Bignon EA, Muena NA, Manuguerra J-C, Jangra RK, et al. The Hantavirus Surface Glycoprotein Lattice and Its Fusion Control Mechanism. Cell. 2020 Sep 15;183(2):442–56.

27. Evans R, O’Neill M, Pritzel A, Antropova N, Senior A, Green T, et al. Protein complex prediction with AlphaFold-Multimer. bioRxiv. 2021 Oct 4;2021.10.04.463034.

28. Guardado-Calvo P, Rey FA. The Viral Class II Membrane Fusion Machinery: Divergent Evolution from an Ancestral Heterodimer. Viruses. 2021 Nov 1;13(12).

29. Lappin DF, Nakitare GW, Palfreyman JW, Elliott RM. Localization of Bunyamwera bunyavirus G1 glycoprotein to the Golgi requires association with G2 but not with NSm. J Virol. 1994 Dec 1;75(Pt 12):3441–51.

30. Shi X, Brauburger K, Elliott RM. Role of N-linked glycans on bunyamwera virus glycoproteins in intracellular trafficking, protein folding, and virus infectivity. J Virol. 2005;79(21):13725–34.

31. Guardado-Calvo P, Bignon EA, Stettner E, Jeffers SA, Pérez-Vargas J, Pehau-Arnaudet G, et al. Mechanistic Insight into Bunyavirus-Induced Membrane Fusion from Structure-Function Analyses of the Hantavirus Envelope Glycoprotein Gc. Kuhn RJ, editor. PLOS Pathog. 2016 Oct 26;12(10):e1005813.

32. Halldorsson S, Li S, Li M, Harlos K, Bowden TA, Huiskonen JT. Shielding and activation of a viral membrane fusion protein. Nat Commun. 2018 Dec 24;9(1):349.

33. Bitto D, Halldorsson S, Caputo A, Huiskonen JT. Low pH and Anionic Lipid-dependent Fusion of Uukuniemi Phlebovirus to Liposomes. J Biol Chem. 2016 Mar 18;291(12):6412–22.

34. Overby A, Pettersson R, Grünewald K, Huiskonen J. Insights into bunyavirus architecture from electron cryotomography of Uukuniemi virus. Proc Natl Acad Sci U S A. 2008 Feb 19;105(7):2375–9.

35. Freiberg A, Sherman M, Morais M, Holbrook M, Watowich S. Three-dimensional organization of Rift Valley fever virus revealed by cryoelectron tomography. J Virol. 2008 Nov 1;82(21):10341–8.

36. Sherman M, Freiberg A, Holbrook M, Watowich S. Single-particle cryo-electron microscopy of Rift Valley fever virus. Virology. 2009 Apr 25;387(1):11–5.

37. Huiskonen J, Overby A, Weber F, Grünewald K. Electron cryo-microscopy and single-particle averaging of Rift Valley fever virus: evidence for GN-GC glycoprotein heterodimers. J Virol. 2009 Apr 15;83(8):3762–9.

38. Voss JE, Vaney MC, Duquerroy S, Vonrhein C, Girard-Blanc C, Crublet E, et al. Glycoprotein organization of Chikungunya virus particles revealed by X-ray crystallography. Nature. 2010 Dec 2;468(7324):709–12.

39. Guardado-Calvo P, Rey FA. The Envelope Proteins of the Bunyavirales. Adv Virus Res. 2017 Jan 1;98:83–118.

40. Guardado-Calvo P, Rey FA. The surface glycoproteins of hantaviruses. Vol. 50, Current Opinion in Virology. Elsevier; 2021. p. 87–94.

41. Bignon EA, Albornoz A, Guardado-Calvo P, Rey FA, Tischler ND. Molecular organization and dynamics of the fusion protein Gc at the hantavirus surface. Elife. 2019 Jun 10;8:e46028.

42. Rissanen I, Stass R, Zeltina A, Li S, Hepojoki J, Harlos K, et al. Structural transitions of the conserved and metastable hantaviral glycoprotein envelope. J Virol. 2017;91(21):378–417.

43. Hofmann H, Li X, Zhang X, Liu W, Kuhl A, Kaup F, et al. Severe Fever with Thrombocytopenia Virus Glycoproteins Are Targeted by Neutralizing Antibodies and Can Use DC-SIGN as a Receptor for pH-Dependent Entry into Human and Animal Cell Lines. J Virol. 2013 Apr 15;87(8):4384–94.

44. Sabella E, Pierro R, Panattoni A, Materazzi A, Vergine M, De Bellis L, et al. Effects of modulation of potassium channels in tobacco mosaic virus elimination. Physiol Mol Plant Pathol. 2018 Apr 1;102:180–4.

45. Pearson H, Todd EJAA, Ahrends M, Hover SE, Whitehouse A, Stacey M, et al. TMEM16A/ANO1 calcium-activated chloride channel as a novel target for the treatment of human respiratory syncytial virus infection. Thorax. 2021 Jan 1;76(1):64–72.

46. Hopkins FR, Álvarez-Rodrígues B, Heath GR, Panayi K, Hover S, Edwards TA, et al. The structure of a native orthobunyavirus ribonucleoprotein reveals a key role for viral RNA in maintaining its helical architecture. bioRxiv. 2021;

47. Zhang K. Gctf: Real-time CTF determination and correction. J Struct Biol. 2016 Jan 1;193(1):1–12.

48. Xiong Q, Morphew MK, Schwartz CL, Hoenger AH, Mastronarde DN. CTF determination and correction for low dose tomographic tilt series. J Struct Biol. 2009 Dec;168(3):378–87.

49. Mastronarde DN, Held SR. Automated tilt series alignment and tomographic reconstruction in IMOD. J Struct Biol. 2017 Feb 1;197(2):102–13.

50. Nicastro D, Schwartz CL, Pierson J, Gaudette R, Porter ME, Richard MJ. The molecular architecture of axonemes revealed by cryoelectron tomography. Science (80-). 2006 Aug 18;313(5789):944–8.

51. Heymann JB, Belnap DM. Bsoft: image processing and molecular modeling for electron microscopy. J Struct Biol. 2007 Jan;157(1):3–18.

52. Pettersen EF, Goddard TD, Huang CC, Couch GS, Greenblatt DM, Meng EC, et al. UCSF Chimera--a visualization system for exploratory research and analysis. J Comput Chem. 2004 Oct;25(13):1605–12.

53. Pettersen EF, Goddard TD, Huang CC, Meng EC, Couch GS, Croll TI, et al. UCSF ChimeraX: Structure visualization for researchers, educators, and developers. Protein Sci. 2021 Jan 1;30(1):70–82.

54. Scott CCC, Gruenberg J. Ion flux and the function of endosomes and lysosomes: pH is just the start. Bioessays. 2011 Feb;33(2):103–10.

